# Combining 3D-multispectral and hyperspectral imaging to identify environmental stress treatments imposed during plant growth

**DOI:** 10.64898/2026.08.28.747774

**Authors:** Frederike Stock, Saswat Panda, Richard Poiré, Timothy Brown, Ayesha Akram, Liang Zheng, Huan Lei, Ruyi Zha, Mingrui Zhao, Sebastien Isabelle, Michèle Martel, Marc-André Comeau, Louis-Philippe Hamel, Pierre-Olivier Lavoie, Marc André D’Aoust, Hannah Reithinger, Pooja Saxena, Eric A. Stone, Hongdong Li, Danielle A. Way, Owen K. Atkin

## Abstract

Non-invasive, high-throughput phenotyping tools are needed that can identify environmental effects on plant structure and function to diagnose factors responsible for reduced growth in commercial and non-commercial settings. In this study, we explored whether the integration of 3D-multispectral (3D) and 2D-hyperspectral imaging (HSI), aided by machine learning (ML), could be used to identify environmental stress treatments imposed during plant growth. Controlled environment-grown *Nicotiana Benthamiana* plants were subjected to a range of abiotic treatments – including different growth irradiances, heat treatment and drought stress – with the treatments resulting in differences in shoot height, biomass, leaf area and spectral reflectance. ML models were trained to identify these treatments using morphological and spectral traits measured at 27, 29, 31, and 34 days after sowing (DAS). A 3D-multispectral scanner was used to obtain information on plant height, biomass, and leaf area. A visible and near-infrared (VNIR) HSI camera provided detailed spectral information for deriving spectral indices including the Normalised Difference Vegetation Index (NDVI), Photochemical Reflectance Index (PRI) and Normalized Difference Red Edge (NDRE). Manual measurements provided baseline comparative data.

The 3D-multispectral scanner reliably estimated above-ground traits, with high correlations between manual and scanner-derived measurements. The ML models accurately differentiated among environmental stress treatments, with the fused 3D+HSI model achieving the best overall predictive performance across all evaluated metrics compared with models based on either imaging modality alone.

Results demonstrated the effectiveness of combining 3D-multispectral and 2D-HSI data with ML analyses for non-destructive, high-throughput phenotyping. The integration of these techniques enabled non-destructive, high-throughput identification of environmental stress treatments imposed during plant growth.

## Introduction

In nature, plants are exposed to ever-changing abiotic environments, with sustained variations in light, temperature and water availability having profound effects on plant growth and performance (Lambers and Oliveira 2019). Changes of the abiotic environment often result in modifications in canopy structure (Nobel and Long 1985) and in modulation of the morphological, anatomical, metabolic and chemical composition of leaves (Atkin, Loveys et al. 2005, Wright, Reich et al. 2006, Poorter, Niinemets et al. 2009, Nicotra, Atkin et al. 2010, Asao, Hayes et al. 2020, Wang, Townsend et al. 2022). For example, plants grown in high light typically exhibit thicker leaves, leading to higher concentrations of photosynthetic pigments per unit leaf area than their low-light-grown counterparts (Evans and Poorter 2001, Terashima, Hanba et al. 2005, Hoshino, Yoshida et al. 2019). By contrast, leaves developed under warmer temperatures are typically thinner than their colder-grown counterparts (Poorter, Niinemets et al. 2009), with lower leaf nitrogen (N) concentrations (Ordonez, Bodegom et al. 2009) and protein content (Scafaro, Xiang et al. 2017). Under drought, leaves often exhibit reduced stomatal conductance (Buckley 2019), accumulation of metabolites that enable osmotic adjustment (Mukarram, Choudhary et al. 2021) and altered pigment composition (Gallé and Feller 2007) – changes that influence the ability of plants to photosynthesise and accumulate biomass. Light quality can also alter relative investment in different pigment types (i.e. chlorophylls, carotenoids and flavonoids), with consequences for photosynthetic efficiency (Demmig-Adams and Adams III 1992). Noting these observations, the structural and chemical composition of leaves and whole canopies reflect the environmental conditions under which plants have grown. Yet this information has rarely been used to investigate environmental conditions experienced by herbaceous plants.

Understanding the environmental conditions to which plants have been exposed during growth is important for a range of industries. For example, reduced plant performance caused by environmental stress can significantly reduce yields in horticulture and in commercial plant production (Lobell and Gourdji 2012, Francini and Sebastiani 2019); however, the cause of these reduced performances can be difficult to establish. Likewise, in broadacre cropping, plants are frequently exposed to a range of environmental stresses simultaneously (e.g. drought, heat, nutrient limitation, disease), but it can be difficult to know, *in situ*, the degree to which each of these stresses are present, and which had the strongest impact on the observed phenotypes (Galieni, D’Ascenzo et al. 2021). Thus, there is a need to develop better tools that pinpoint the specific causes that result in reduced plant performance, as well as to detect these impacts rapidly so they can be controlled before yield reduction occurs (Grieve, Duckett et al. 2019). As the field of high-throughput plant phenotyping has matured over the past decade, there have been significant improvements in our ability to measure plant traits and quantify plant responses to applied treatments. This maturation has occurred through gradual improvements in phenotyping hardware, data management, and analysis software. Initially, top-down 2D imaging using conventional colour (RGB) cameras was developed for phenotyping model plants with a flat rosette, such as *Arabidopsis thaliana* (Green, Appel et al. 2012, Vasseur, Exposito-Alonso et al. 2018). However, 2D imaging is ineffective for most larger plants, crops and plants with narrow leaves. This technology gap led to the development of modern 3D imaging techniques and hardware platforms that can provide growth rate and leaf area data (Li, Guo et al. 2020, Liu, Bruning et al. 2020). More recently, hyperspectral imaging (HSI) has been used as a plant phenomics tool to address a broad range of questions and systems. HSI can be used to predict a range of plant traits, including physiological functions such as photosynthesis, respiration and water content of plant tissues (Silva-Perez, Molero et al. 2018, Coast, Shah et al. 2019), metabolite concentrations (Vergara-Diaz, Vatter et al. 2020) and responses to abiotic stressors, including drought and heat (Behmann, Steinrücken et al. 2014, Melandri, Thorp et al. 2021, Faqeerzada, Park et al. 2023).

While HSI is a powerful tool, one of the limitations of most HSI platforms is that data are captured using line scanning. Unlike a conventional camera, where the lens captures the entire image at once, a line scanner captures a single strip of the image, one line of pixels at a time, as the scanner travels over the plant. This method enables the camera to capture an image with hundreds to thousands of spectral bands, but the resulting image is 2D, limiting the application of HSI to complex canopies and providing insights only into the leaves at the upper surface of the scan. While some 3D canopy imaging systems integrate multispectral lighting to enable measurements of spectral indices (such as the Normalised Difference Vegetation Index (NDVI) which quantifies vegetation greenness and photosynthetic activity; Table 1) mapped onto 3D plant architecture (Liu, Bruning et al. 2020), no currently available systems integrate hyperspectral data with 3D imaging due to the complexities associated with capturing, fusing and analysing these data types.

**Table 1.** Hyperspectral indices used as inputs for the ML models in this paper.

| Index | Description | Formula | Reference |
| --- | --- | --- | --- |
| NDVI | Normalised Difference Vegetation Index | $\frac{Ref. \text{ at } 800 \text{ nm} - Ref. \text{ at } 680 \text{ nm}}{Ref. \text{ at } 800 \text{ nm} + Ref. \text{ at } 680 \text{ nm}}$ | (Lowe, Harrison et al. 2017) |
| NPCI | Normalised Pigment Chlorophyll Index | $\frac{Ref. \text{ at } 680 \text{ nm} - Ref. \text{ at } 430 \text{ nm}}{Ref. \text{ at } 680 \text{ nm} + Ref. \text{ at } 430 \text{ nm}}$ | (Lowe, Harrison et al. 2017) |
| PSRI | Plant Senescence Reflectance Index | $\frac{Ref. \text{ at } 680 \text{ nm} - Ref. \text{ at } 531 \text{ nm}}{Ref. \text{ at } 800 \text{ nm}}$ | (Lowe, Harrison et al. 2017) |
| PRI | Photochemical Reflectance Index | $\frac{Ref. \text{ at } 531 \text{ nm} - Ref. \text{ at } 570 \text{ nm}}{Ref. \text{ at } 531 \text{ nm} + Ref. \text{ at } 570 \text{ nm}}$ | (Garbulsky, Peñuelas et al. 2011) |
| SR | Simple Ratio | $\frac{Ref. \text{ at } 800 \text{ nm}}{Ref. \text{ at } 680 \text{ nm}}$ | (Lowe, Harrison et al. 2017) |
| SIPI | Structure-Insensitive Pigment Index | $\frac{Ref. \text{ at } 800 \text{ nm} - Ref. \text{ at } 445 \text{ nm}}{Ref. \text{ at } 800 \text{ nm} - Ref. \text{ at } 680 \text{ nm}}$ | (Lowe, Harrison et al. 2017) |
| RENDVI | Red edge Normalised | $\frac{Ref. \text{ at } 750 \text{ nm} - Ref. \text{ at } 705 \text{ nm}}{Ref. \text{ at } 750 \text{ nm} + Ref. \text{ at } 705 \text{ nm}}$ | (Lowe, Harrison et al. 2017) |

|  | Difference Vegetation Index |  |  |
| --- | --- | --- | --- |
| NPQI | Normalised Phaeophytinization Index | $\frac{Ref. \text{ at } 415 \text{ nm} - Ref. \text{ at } 430 \text{ nm}}{Ref. \text{ at } 415 \text{ nm} + Ref. \text{ at } 430 \text{ nm}}$ | Modified from Lowe, Harrison et al. (2017), replacing 435 nm with 430 nm to avoid reflectance loss because of interference of external lights. |
| NDRE | Normalised Difference Red Edge | $\frac{Ref. \text{ at } 800 \text{ nm} - Ref. \text{ at } 720 \text{ nm}}{Ref. \text{ at } 800 \text{ nm} + Ref. \text{ at } 720 \text{ nm}}$ | (Boiarskii and Hasegawa 2019) |
| CCCI | Canopy Chlorophyll Content Index | $\frac{NDRE - NDRE_{min}}{NDRE_{max} - NDRE_{min}}$ | (Cammarano, Fitzgerald et al. 2011) |
| CRI2 | Carotenoid Reflectance Index 2 | $\frac{1}{Ref. \text{ at } 510 \text{ nm}} - \frac{1}{Ref. \text{ at } 700 \text{ nm}}$ | (Wong Man Sing, Tang Hon Wai et al. 2020) |

In this study, we use established 3D-multispectral and 2D-hyperspectral phenotyping tools to test a phenomics proof-of-concept: whether the structural and spectral phenotype of a plant can be used to infer the environmental conditions it has experienced throughout development. The innovation of the work lies in applying multimodal phenotyping to plant environmental detection and in evaluating whether fusing canopy-structural and leaf-spectral traits improves stress classification relative to either modality alone. Our working assumptions were that:

1. ML algorithms trained using 2D-hyperspectral imaging (HSI) data would be superior to models trained on 3D-multispectral data at classifying stress environments that altered a wide range of leaf chemical and structural traits.
2. ML algorithms trained against data from a 3D-multispectral scanner would be capable of classifying environmental conditions that impact canopy structure and/or simple spectral traits linked to leaf greenness (e.g. chlorophyll content) and photosynthetic efficiency.
3. The best ML classifications of plant environmental conditions would be achieved by combining information on canopy structure from a 3D-multispectral scanner and leaf chemical composition information derived from 2D-HSI.

## Materials and Methods

### Plant growth conditions

Seeds of *N. benthamiana* were sown onto rooting plugs (Eazy Plug Rooting Cubes, Eazy Plug, Goirle, The Netherlands), and germination was allowed to proceed for three days in controlled conditions (Adaptis A1000, Conviron, Winnipeg, Canada) at 28/28°C (day/night temperatures), a photoperiod of 16 h :8 h (day/night), and 80% relative humidity (RH) at low light (11 µmol photons m^−2^ s^−1^). During the germination phase, plugs were kept wet using a layer of water in the plug tray. After germination, light intensity in the chamber was increased to 160 µmol photons m^−2^ s^−1^ until nine days after sowing (DAS,) and plugs were sub-irrigated with water daily. From 9-15 DAS, temperature in the growth chamber was decreased to 26/26°C (day/night temperatures), and relative humidity was set to 65%. Seedlings were sub-irrigated daily with a nutrient solution adjusted to an electric conductivity (EC) of 1.0 mS cm^−1^ (Canna Terra Vega, CANNA Australasia, Subiaco, Australia). At 15 DAS, seedlings were transplanted into 500 mL pots with nutrient-rich soil mix (Canna Terra Professional, CANNA Australasia, Subiaco, Australia) and moved into large, controlled environment walk-in growth chambers (Growth Capsule, Photon Systems Instruments, Drásov, Czech Republic). All walk-in growth chambers were equipped with multispectral LEDs with white, red and far-red channels. Between 15 and 25 DAS, plants were sub-irrigated every other day with a nutrient solution of 1.0 mS cm^−1^ until 19 DAS and 1.6 mS cm^−1^ until 25 DAS. From 26 DAS plants were sub-irrigated daily with a nutrient solution adjusted to 2.6 mS cm^−1^.

### Validation of 3D-multispectral imaging using destructive measurements

Morphological and spectral traits of *N. benthamiana* were measured using a 3D-multispectral scanner (Phenospex PlantEye F500 Dualscan, Phenospex, Heerlen, The Netherlands). The 3D-multispectral scanner generates point clouds from individual plants from which several traits are extracted: plant height, digital biomass, 3D leaf area, average canopy greenness, and hue (the angle of the colour on the RGB colour circle). Digital biomass is a volumetric trait (mm^3^) and is calculated by the scanner as a product of plant height (mm) and 3D leaf area (mm^2^). It is not a direct measure of biomass but is expected to show high correlation with actual canopy biomass. Equipped with four bands - red, green, blue and near-infra-red, several vegetation indices could also be calculated, namely the Normalised Difference Vegetation Index (NDVI), the Normalised Pigments Chlorophyll ratio Index (NPCI), and the Plant Senescence Reflectance Index (PSRI). The complete list of scanner-derived traits used in the 3D-multispectral model is provided in Appendix 5.

To validate the 3D scan data, plant height, leaf area and digital biomass of *N. benthamiana* were subsequently compared to manual measurements of plant height, total leaf area per plant, and shoot fresh mass. For the comparison, plants of different developmental stages (21 to 34 DAS) were used. Measurements of plant height were manually assessed by measuring the distance between the base of the stem and the top of the shoot with a ruler. Destructive measurements of total leaf area per plant were taken by dissecting leaves from each plant, lying them flat on a table and taking a photo. Leaf area was then calculated from the photos using the ImageJ Software (Schindelin, Arganda-Carreras et al. 2012). Shoot fresh mass was determined by harvesting all leaves from a plant, including stems, and measuring their total mass using a balance. Correlations between predicted traits from the 3D scanner and manual measurements were determined using linear regression. Coefficient of determination (R^2^), *p*-values, slope and intercept values were calculated using the scipy.stats package from Python (Virtanen, Gommers et al. 2020).

### Exposure of plants to a range of environmental conditions

Four separate experiments were used to monitor changes in *N. benthamiana* phenotypes under heat, drought, irradiance, or selected treatment combinations (Drought + Light; Heat + Drought) from 15 DAS until harvest (Table 2). Each experiment comprised 20-52 plants, with 10-32 plants per treatment (Table 2). The photoperiod for all environmental conditions was 16h:8h (day/night). Abiotic treatments were applied on a subset of plants from 15 DAS by adjusting the relevant environmental conditions in the growth chamber. For control treatments (CTRL), plants were grown at 25/22.5 °C (day/night), 50% RH and a light intensity of 300 µmol photons m^−2^ s^−1^. For the two light treatments, irradiance was increased to either 450 µmol photons m^−2^ s^−1^ (Light) or 600 µmol photons m^−2^ s^−1^ (Light) by adjusting the white light channel. This adjustment also led to an increase in blue light while red and far-red light intensities remained constant. Consequently, changes in light intensity were accompanied by slight changes in the spectral composition (see Appendix 1). Drought treatment commenced 15 DAS and lasted until harvest. Soil moisture in the pots was monitored gravimetrically by weighing every 1-2 days, with drought treated plants maintained at 40% field capacity, while control plants were maintained at 70-80% field capacity. For the heat treatment (HT), temperatures in the growth space were increased to 30/22°C (day/night) in Experiment 1 and to 30/27.5°C in Experiment 4. Although both regimes represented elevated-temperature treatments, their different night temperatures could produce non-equivalent phenotypic responses. They were grouped into a single broad Heat category for model training, and a sensitivity analysis comparing their within-experiment, control-relative phenotypic signatures was performed to evaluate this grouping. Two combinatorial treatments were also used: a drought treatment was combined with a high light treatment in Experiment 3 (Drought + Light_450_) and a heat treatment was combined with a drought treatment in Experiment 4 (Heat + Drought) (Table 2). For each experiment, 2 or 3 controlled environment growth chambers (Photon System Instruments, Growth Capsules) were used to apply the treatments: for Experiment 1, two growth rooms were used (one for control and one for heat); for Experiment 2, three growth rooms were used (control, Light_450_ and Light_600_); for Experiment 3, two growth rooms were used (control + drought, Light_450_ + drought); and for Experiment 4, two growth rooms were used (control + drought, heat + drought). Because control phenotypes varied modestly across experiments and treatment assignment was partially associated with batch and growth chamber, we evaluated a fold-wise batch-control-standardised version of the primary leave-one-out analysis. To avoid information leakage, normalization parameters were estimated using training controls only within each fold (Varma and Simon 2006; Cawley and Talbot 2010).

**Table 2.** Overview of the four experiments performed in this study. Each experiment contained a control treatment and one or more environmental treatments. The number of plants used for each treatment is shown, as are the number of plants scanned using 3D structure and hyperspectral reflectance.

| Experiment | Total # of plants | Treatments | # plants per treatment | # plants with 3D & hyperspectral scans |
| --- | --- | --- | --- | --- |
| 1 | 20 | Control<br>Heat | 10<br>10 | 9<br>9 |
| 2 | 48 | Control<br>Light<sub>450</sub><br>Light<sub>600</sub> | 16<br>16<br>16 | 10<br>10<br>10 |
| 3 | 45 | Control<br>Drought<br>Drought + Light<sub>450</sub> | 10<br>15<br>20 | 5<br>10<br>15 |
| 4 | 52 | Control<br>Drought<br>Heat<br>Heat + Drought | 10<br>10<br>12<br>20 | 4<br>5<br>6<br>15 |

### Plant phenotyping

During each experiment, plants were scanned 2 or 3 times per week using the 3D-multispectral scanner and a 2D VNIR hyperspectral camera (400-900 nm, 500 * 385 pixels, Photon Systems Instruments, Drásov, Czech Republic). For 18 plants from the Control and Heat treatments of Experiment 1, hyperspectral scanning data was unavailable at 29 DAS. These missing data points were linearly interpolated before training the ML model. For further details, please refer to Appendix 2. Destructive harvesting of three plants per treatment was performed at the end of each experiment (34 DAS) and statistical analyses of the data were performed using GraphPad Prism (version 10.1.0 for Windows, GraphPad Software, Boston, MA, USA). For Experiment 1, differences in means presented in Figure 2 were tested using an unpaired Student’s t-test. For Experiments 2-4, differences in means were analysed using one-way ANOVA followed by Dunnett’s multiple comparisons test.

### Pre-processing of the hyperspectral data

Several hyperspectral indices were calculated from the hyperspectral scanner data for use by a machine learning (ML) model (Table 1). These indices were selected because they are biologically interpretable and capture pigment composition, photosynthetic light-use efficiency, chlorophyll status and senescence-related spectral responses. Calculation of indices required pre-processing of hyperspectral images, including white-dark calibration, index computation, and background removal. White-dark calibration corrects the spectral response of the hyperspectral system for background and sensor noise. Calibration was achieved by using the white reference (generated by scanning a Spectralon panel) and a dark reference (an image taken with closed shutter) to correct hyperspectral images of *N. benthamiana* using the formula ((sample – dark) / (white – dark)) in Equation (1) in Lu and Chen (1999). For index computation, the required wavelengths were identified in the image output and selected indices were calculated for each pixel in the hyperspectral image. We intentionally used a limited set of established, biologically interpretable hyperspectral indices rather than the full raw spectrum. This choice was made to reduce the risk of overfitting, because the number of raw spectral bands would be high relative to the 108 plant-level samples available for modelling (Hughes 1968). A second reason for using the indices was to determine whether a small number of informative wavelength combinations could improve classification accuracy, which may inform future reduced-band or multispectral sensor configurations without requiring full-spectrum HSI acquisition.

After calculating the selected indices, pixels that were out of range were clipped to mitigate the effects of extreme values. Finally, the background was removed by separating the plant canopy from the image background. To achieve this, a mask was generated by applying a combination of green colour detection (MathWorks documentation) and Otsu’s thresholding (Otsu 1979) on the reference colour image generated by the hyperspectral camera. Subsequently, the average value of all pixels of the plant was computed, resulting in an averaged index per plant. This single scalar of each plant was then used as input for the ML model (for details refer to Appendix 3).

### Development of ML models

Identifying plant stress, as intended in this study, is a classification problem in the field of ML (Mohanty, Hughes et al. 2016). Considering the limited number of samples in each treatment (Table 3), the popular ML model XGBoost (Chen and Guestrin 2016) was used, because of its interpretability and ability to learn accurately from small-sized datasets. The same approach would not have been possible with deep learning approaches (Shahinfar, Meek et al. 2020). Given the constraints of the dataset and the objective to minimise overfitting while maintaining robust model performance, the XGBoost classifier was configured with a maximum tree depth of four (4) and a learning rate of 0.1, for the primary leave-one-out testing analysis. These settings were chosen to constrain model complexity and avoid overly deep trees in a dataset with a limited number of plant-level samples. We did not optimise the model parameters or perform comprehensive feature selection within the primary leave-one-out testing procedure, because carrying out model selection and testing on the same small dataset can lead to selection bias and optimistic performance estimates unless model selection is nested within the validation procedure (Varma and Simon 2006; Cawley and Talbot 2010). Although hyperparameter tuning is typically recommended using grid search or random search (Pedregosa, Varoquaux et al. 2011), this was not feasible in this case due to the lack of a separate test set. Therefore, leave-one-out testing (for details refer to Appendix 4), a variant of cross-validation, was selected as the primary testing procedure to maximise data utilisation. In each iteration, one plant-level sample was held out for testing, and the model was trained on the remaining plants. This ensured that the test plant was not used during model fitting and avoided leakage of plant-specific information between training and testing. The final performance was calculated by aggregating the predictions across all 108 held-out plant-level tests. The random state was set to a fixed value to guarantee the reproducibility of the model training process. In this paper, feature weighting refers to XGBoost normalized mean-gain feature importance (Chen and Guestrin 2016). Weightings were calculated separately in each LOOCV fold. For each fold, the weightings for DAS-specific columns belonging to the same underlying trait were summed, expressed as percentages, and then averaged across folds. These model-derived weightings describe predictive contribution within the fitted models; they do not establish causal physiological importance, and correlated predictors may share or redistribute weighting.

**Table 3.** Dataset used to train the ML models and for the identification of single and combined stress treatments. Each sample represents a single plant measured on multiple DAS (27, 29, 31 and 34) as they developed under the treatment conditions. For more details about the abiotic treatments, refer to plant growth conditions in the Materials and Methods section.

| Treatment | Number of samples |
| --- | --- |
| Control | 28 |
| Heat | 15 |
| Drought | 15 |
| Light<sub>450</sub> | 10 |
| Light<sub>600</sub> | 10 |
| Drought +<br>Light<sub>450</sub> | 15 |
| Drought + Heat | 15 |
| **Total samples** | **108** |

For statistical comparison of model performance, paired tests were used because the three models were evaluated on the same 108 plants. Cochran’s Q test was used for the overall comparison among the three paired model outcomes, followed by pairwise exact McNemar tests with Holm correction for multiple comparisons and bootstrap resampling to estimate confidence intervals for paired accuracy differences (Cochran 1950; McNemar 1947; Holm 1979; Efron 1979). Temporal sensitivity was examined by repeating the fused 3D+HSI analysis using the same model configuration with (i) all 15 traits measured at 27, 29, 31 and 34 DAS (60 trait-by-time features), (ii) the 15 traits measured at 31 DAS only, and (iii) the 15 traits measured at 34 DAS only. Complete-history and single-date models were compared using paired exact McNemar tests with Holm correction and paired bootstrap confidence intervals. Batch and chamber sensitivity was evaluated by comparing the primary raw-feature LOOCV with a fold-wise batch-control-standardised LOOCV. Normalization parameters were estimated using training controls only within each fold to avoid information leakage. Paired outcomes were compared using Cochran’s Q test, an exact McNemar test, and a paired bootstrap confidence interval for the accuracy difference. Heat-regime sensitivity was evaluated by comparing the within-experiment, control-relative signatures of the two heat regimes across the 60 fused 3D+HSI features. Comparisons used Pearson and Spearman correlations, cosine similarity, and effect-direction agreement. Feature-level differences were assessed using Welch and Mann–Whitney tests with false-discovery-rate correction (Welch 1947; Mann and Whitney 1947; Benjamini and Hochberg 1995). Model confidence was assessed using the maximum predicted-class probability for each of the 108 out-of-fold predictions. Confidence scores for correct and incorrect classifications were compared using a two-sided Mann–Whitney U test. Misclassified plants were also summarized by experiment based on the batch identifier in each plant ID. Maximum predicted-class probabilities were interpreted as relative model-confidence scores rather than formally calibrated probabilities (Niculescu-Mizil and Caruana 2005). To evaluate parameter sensitivity and classifier choice independently of the primary leave-one-out analysis, we performed an additional representative 75/25 train/test split analysis. Grid-search tuning was performed using the training set, and the selected models were then evaluated on the held-out test set of 27 plants. This analysis also compared XGBoost with Random Forest, Linear SVM and Logistic Regression. It was not used to optimise or alter the parameters used in the primary plant-level leave-one-out testing experiment. For details, refer to Appendix 7.

## Results

### Validation of morphological plant traits measured by the 3D scanner

To ensure that the 3D-multispectral scanner reliably estimated morphological traits of *N. benthamiana*, plant height, leaf area and digital biomass predictions by the scanner were compared to empirical measurements of these traits (Figure 1 **Error! Reference source not found**.). In the three cases, significant correlation levels (R² values > 0.9; *p*-value < 0.0001) were denoted when comparing readings of the 3D-multispectral scanner with empirical measurements of the corresponding traits. For plant height (Figure 1a), the slope of the relationship between manual measurements and the predicted plant height from the 3D scans was 0.86, indicating that both approaches yielded very similar values. By contrast, the correlation of the 2D and 3D leaf areas yielded a slope of 2.3, indicating that the 3D scanner underestimates the “true” 2D leaf area due to occlusion, especially as the plants grow bigger and the canopy becomes more complex. Despite the underestimation of leaf area, the correlation between both measurements was high (R² value > 0.92) (Figure 1b). Shoot biomass measured by the 3D scanner (digital biomass) is a product of plant height and 3D leaf area and therefore results in a volumetric measurement which is a different metric than traditional aboveground biomass measurements. Digital biomass measurements showed good correlation with destructive shoot biomass measurements (R² values > 0.92) (Figure 1c).

**Figure 1.**
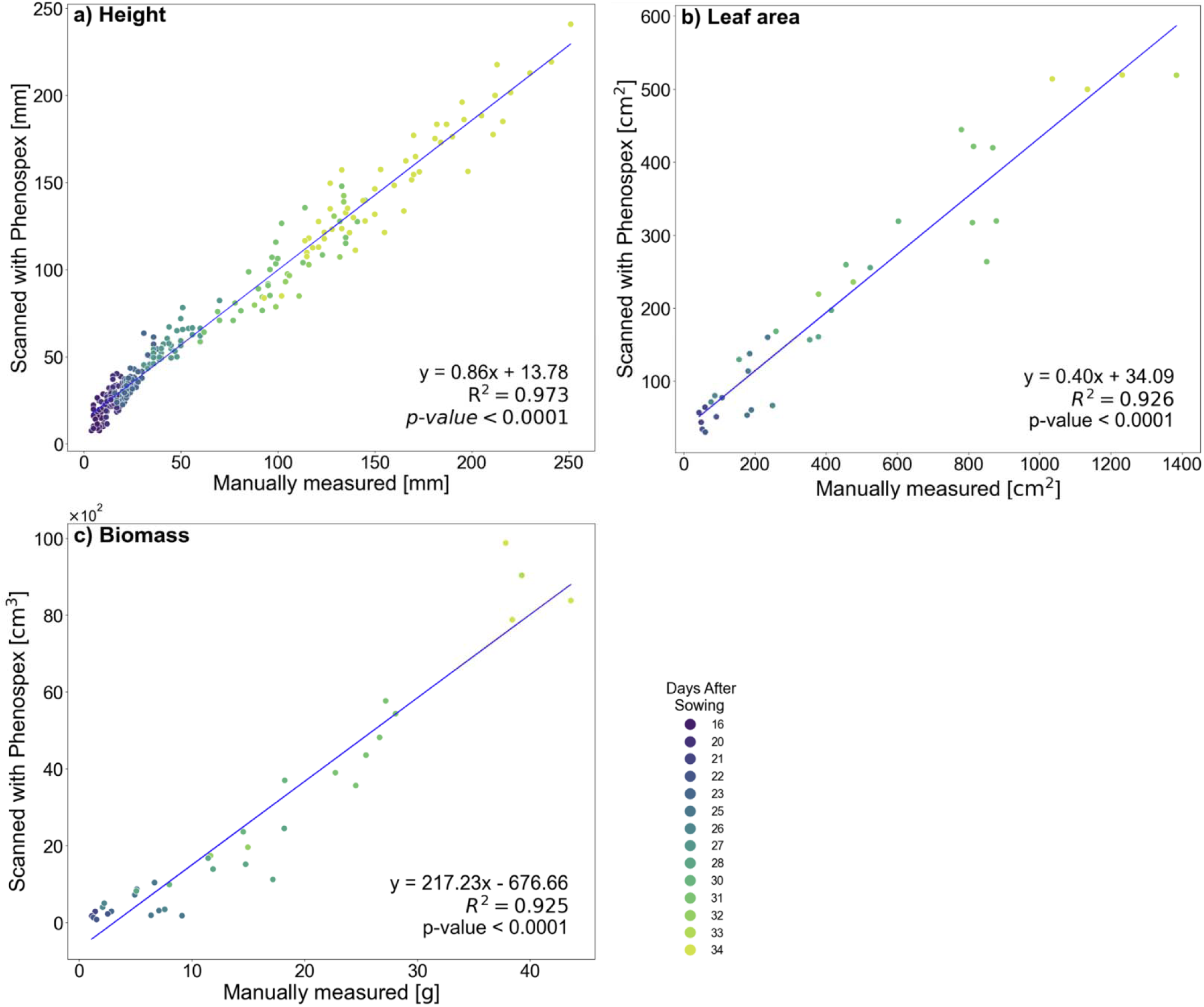
Correlation of plant height (a), leaf area (b), and aboveground biomass (c), as measured by the 3D multispectral scanner (y-axes) compared to the empirical measurements (x-axes). Each data point re resents the data from a single plant at a given DAS (days after sowing). Colours indicate the age of the plant ranging from young plants (16 DAS) marked in purple, to older plants fading to yellow (34 DAS). High levels of correlation were observed for all three comparisons, as indicated with R values > 0.9. (a) n = 272, (b n = 35, and (c) n = 37.

### Plant growth responses to environmental treatments

Having established the reliability of morphological traits from the 3D-multispectral scanner, *N. benthamiana* plants from control and abiotic stress treatments were scanned 2 or 3 times per week with the 3D-multispectral scanner after being transplanted into pots 15 DAS. A subset of the plants was also used for measurements with the hyperspectral scanner on the same days to obtain detailed spectral information on the canopies (Table 2).

Distinct changes were observed in plant height, leaf area and biomass in response to the abiotic treatments across all four experiments, as confirmed by the destructive measurements (Figure 2). For example, the Heat treatment led to an increase in plant height (Figure 2a, d) and leaf area (Figure 2 **Error! Reference source not found**.e, h) in Experiments 1 and 4, but had no measurable impact on shoot biomass (Figure 2i, l). By contrast, morphological effects of the Drought treatment (Experiments 3 and 4) were less pronounced, with no significant impact on plant height or leaf area (Figure 2c, d, g, h), but a decrease in biomass (Figure 2k, l). The Light treatments (Experiments 2 and 3) did not affect plant height nor shoot biomass, but higher light intensities increased leaf area in Experiment 2 (Figure 2f). The two heat regimes showed an overall shared phenotypic pattern, with 43/60 features (trait-by-time combinations; 71.7%) changing in the same direction relative to their respective controls. However, 22/60 features differed significantly between the regimes after FDR correction. We therefore grouped them as a broad elevated-temperature class for model development while recognizing that the regimes were not phenotypically equivalent. The difference in night temperature is retained as a study limitation (Appendix 11).

**Figure 2.**
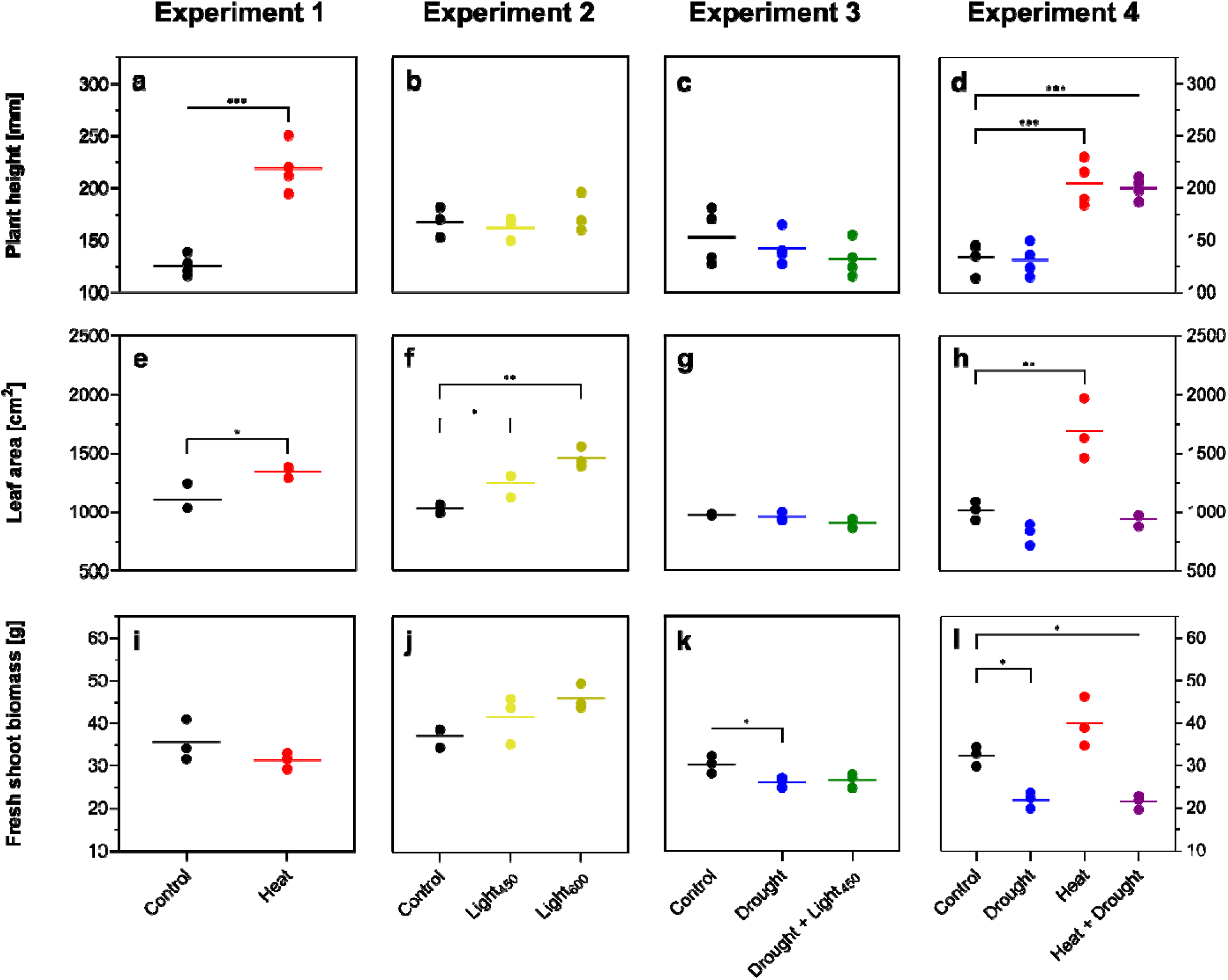
Empirically measured plant height, leaf area and fresh shoot biomass of *N. benthamiana* lants at 34 DAS. Leaf area and biomass were measured destructively. Horizontal lines indicate the mean of each treatment group. Various colors depict the different treatments and asterisks (*) highlight si nificant differences between conditions (* = p-value ≤ 0.05; ** = p-value ≤ 0.01; *** = p-value ≤ 0.001). Graphs b) and e-l) n=3; graphs a), c) and d) n=3.

### Comparison of three ML models

After characterizing plants from the four experiments with both the 3D-multispectral scanner and the hyperspectral camera, the resulting data were used to train three ML models to differentiate among plants grown under each environmental condition (Table 4). One model only used data from the 3D-multispectral scanner as an input (3D-model), one model only used the 2D-HSI data as an input (HSI-model), and the third model used both datasets as an input (3D + HSI-model). For environment treatment identification at 34 DAS, the overall accuracy achieved by the models was the highest for the 3D + HSI-model, followed by the HSI-model and the 3D-model (Table 4). The full trait list and weightings for each model are provided in Appendix 5. Briefly, the 3D model used 12 3D-multispectral traits, the HSI model used 11 hyperspectral indices, and the fused 3D+HSI model used 11 3D-multispectral traits together with 4 HSI-derived indices.

**Table 4.** Comparison of test accuracy (%) at 34 DAS for the 3D-, HSI- and 3D+HSI-models. For confusion matrices, complete trait lists, trait weightings and additional model-performance metrics, refer to Appendices 5 and 6 in the Supplementary Information. Values displayed in bold indicate the highest accuracy for identifying plants grown in each abiotic treatment. For details on the hyperspectral indices, refer to Table 1.

| Treatment | 3D-model (%) | HSI-model (%) | 3D + HSI-model (%) |
| --- | --- | --- | --- |
| Control | 79 | 86 | **89** |
| Heat | **93** | 80 | **93** |
| Drought | 40 | 73 | **80** |
| Light<sub>450</sub> | 40 | 70 | **90** |
| Light<sub>600</sub> | 60 | **80** | **80** |
| Drought + Light<sub>450</sub> | 73 | **100** | 93 |
| Drought + Heat | **100** | 73 | 93 |
| Overall | 72 | 81 | **89** |
| **Most weighted traits** | Digital Biomass, Greenness, Height, PSRI | PRI, NDRE, RENDVI, NPCI | HSI: PRI and NDRE<br>3D: PSRI & Digital Biomass |
| **Accuracy distribution across treatments** | Comparatively least balanced accuracy across treatments. | Comparatively better balancing of accuracies. | Comparatively best balancing of accuracies across treatments. |

Across the 108 plant-level leave-one-out tests, the fused 3D+HSI model achieved the highest overall performance, with an accuracy of 88.9%, balanced accuracy of 88.5% and macro F1-score of 88.0%. This was higher than the HSI-only model, which achieved 81.5% accuracy, 80.3% balanced accuracy and 79.7% macro F1-score, and the 3D model, which achieved 72.2% accuracy, 69.3% balanced accuracy and 68.3% macro F1-score. Because the three models were evaluated on the same plants, paired statistical tests were used to compare model correctness. Cochran’s Q test indicated a significant overall difference among the three paired model outcomes (Q = 11.619, df = 2, p = 0.002999). Pairwise exact McNemar tests with Holm correction showed that the fused 3D+HSI model significantly outperformed the 3D model, whereas the 3D and HSI models did not differ significantly. The fused 3D+HSI model also showed directional evidence of improvement over the HSI-only model, supported by a positive bootstrap accuracy-difference interval, although this comparison did not meet significance under the two-sided Holm-corrected McNemar test. Detailed model-comparison results, including per-class metrics, paired statistical tests and bootstrap accuracy differences, are provided in Appendix 6.

In a separate batch-control sensitivity analysis, the difference in accuracy between the fused model using raw trait measurements and the same model using fold-wise batch-control-standardised trait measurements was not statistically significant. This result suggests that classification performance was not solely driven by batch-specific control differences, although residual confounding by batch and growth chamber cannot be excluded (Appendix 9).

To assess whether the classification performance was specific to XGBoost or reflected separability in the phenotypic data, we also compared XGBoost with Random Forest, Linear SVM and Logistic Regression using a representative 75/25 held-out grid-search analysis. In this supplementary analysis, the fused 3D+HSI feature set achieved high held-out performance across multiple classifier types, supporting the conclusion that multimodal phenotypic data were strongly discriminative rather than the result being unique to XGBoost alone (Appendix 7).

The 3D-model performed best at identifying plants submitted to Heat and Drought + Heat treatments, while the HSI-model most accurately identified plants grown in the Drought+Light_450_ and Light_600_ treatments. The relatively low 3D-model accuracy for Drought and Light450 likely reflects the weaker structural separation of these treatments compared with Heat and Drought+Heat. Drought and Light450 produced less consistent differences in plant height, projected leaf area and digital biomass, making them more difficult to classify using 3D structural traits alone. By contrast, these treatments can alter pigment composition, chlorophyll status and photochemical activity, which are more directly captured by the HSI-derived indices. The 3D+HSI-model enabled the identification of all abiotic treatments with ≥ 80% accuracy, and presented the highest accuracy for the Control, Heat, Drought, Light_450_ and Light_600_ treatments. Based on the confusion matrices (for details, refer to Appendix 5), the 3D model showed the strongest confusion between Drought and Drought+Light450, consistent with the relatively weak structural separation of these treatments. This confusion was reduced in the HSI model and was lowest in the fused 3D+HSI model. In the fused model, the largest remaining confusion was between Light450 and Light600, suggesting that these two irradiance treatments produced partially overlapping phenotypic signatures, particularly when analysed at the whole-plant trait level.

The ML model used different weightings for different input traits to differentiate among the abiotic treatments. Weightings used in all three models are provided in Appendix 5, allowing comparison of which structural and spectral traits contributed most strongly to each classification model. For example, in the 3D+HSI-model, PRI (an index sensitive to photosynthetic light use efficiency) was assigned the maximum weighting, as it effectively differentiates between four treatments (Figure 3a). PRI did not clearly differentiate between the Heat and Light_450_ treatments up to 29 DAS and therefore the ML model incorporated the weighting of Digital Biomass, which shows a clear distinction between these two treatments (Figure 3b). The second-most weighted index was NDRE (a red-edge index sensitive to chlorophyll content), which enabled the models to distinguish Drought and Drought+Light_450_ (Figure 3c). NDRE also differentiated among the four treatments (Figure 3a and 3c) by 31-34 DAS, but did not differentiate between the Drought+Light_450_ and Drought+Heat treatments up to 29 DAS. Noting this, the model therefore added weightings to NPCI to differentiate these two treatments more clearly (Figure 3d).

**Figure 3.**
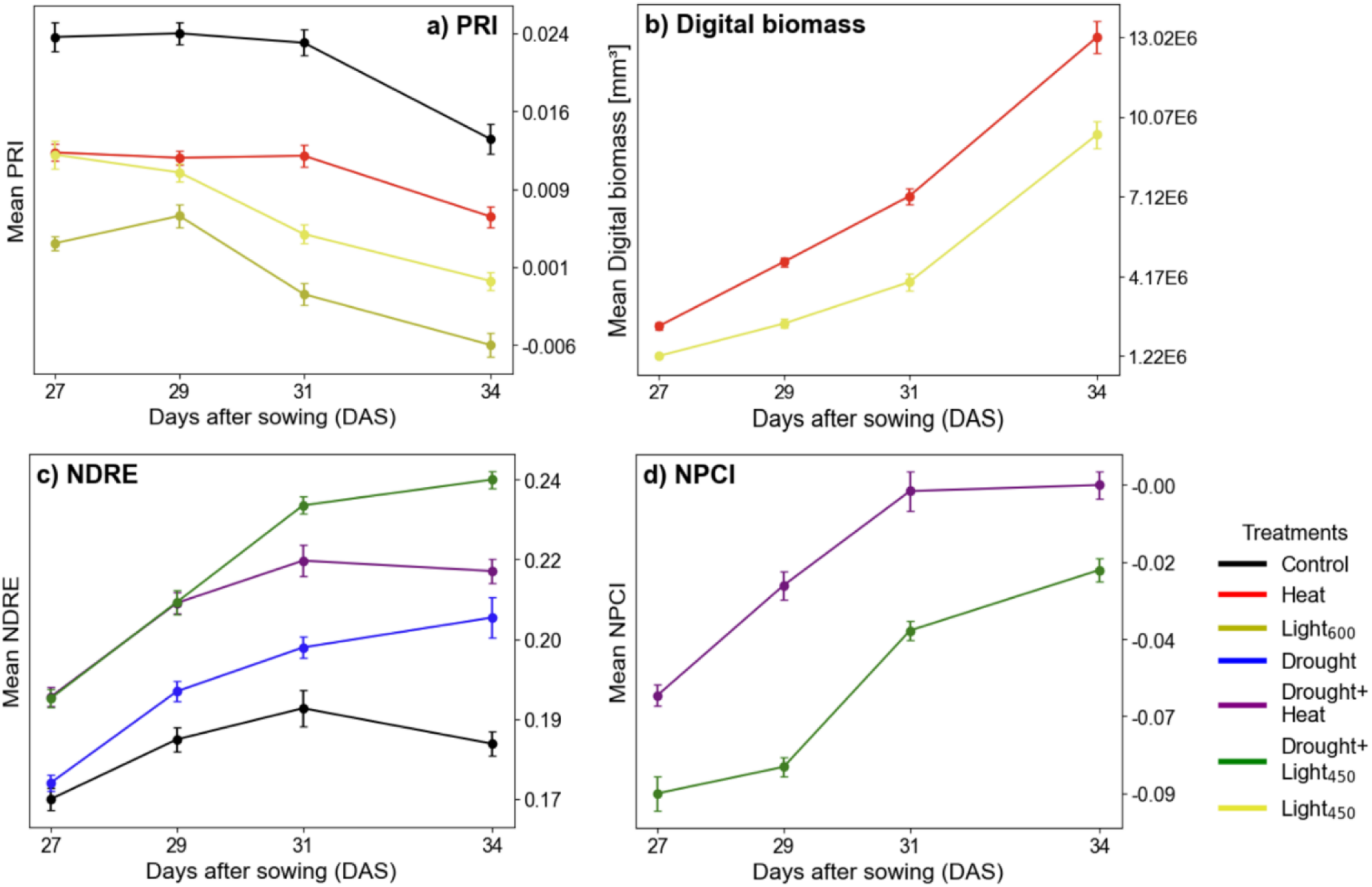
Multi-faceted analysis of plant responses to various abiotic treatments over time, as measured in days after sowing (DAS). Each panel depicts how a specific trait — Photochemical Reflectance Index (PRI), Digital Biomass, Normalized Difference Red Edge (NDRE), and Normalized Pigment Chlorophyll Index (NPCI) — vary within plants under different abiotic treatments. Panel (a) shows how PRI differentiates between four treatments, as indicated at the bottom right. Panel (b) monitors Digital Biomass, which differentiates Heat and Light_450_ more clearly when compared to PRI. Panel (c) highlights how NDRE differentiates between four treatments, and panel (d) depicts NPCI variations, which can more clearly differentiate Drought + Heat and Drought+Light_450_ compared to NDRE. These indices together with other traits (see Appendix 5) were collectively used by the 3D+HSI-model to accurately identify abiotic treatments imposed to the plants. Error bars represent the standard error (SE). The SE was calculated for each treatment group at a given DAS, with sample sizes (n) ranging from 10 to 28 (see Table 3). Various colors depict the different treatments.

To visualise pixel-by-pixel variations among plants under different abiotic treatments, colour mapping was used, with each index employing a different colour-map. In Figure 4a **Error! Reference source not found**., low PRI (i.e. low light-use efficiency) was depicted in red, while high PRI was depicted in blue. An increase in redness (i.e. a decrease in PRI) was observed in plants as light intensity increased from Light_450_ to Light_600_. In Figure 4b **Error! Reference source not found**., NDRE (an index of chlorophyll content) levels are depicted using a yellow (low) to red (high) gradient. The lowest NDRE was observed in the Control plants, whereas Drought plants exhibited higher NDRE, and plants from the Drought+Light_450_ treatment had the highest NDRE among the three abiotic treatments shown in Figure 4b.

**Figure 4.**
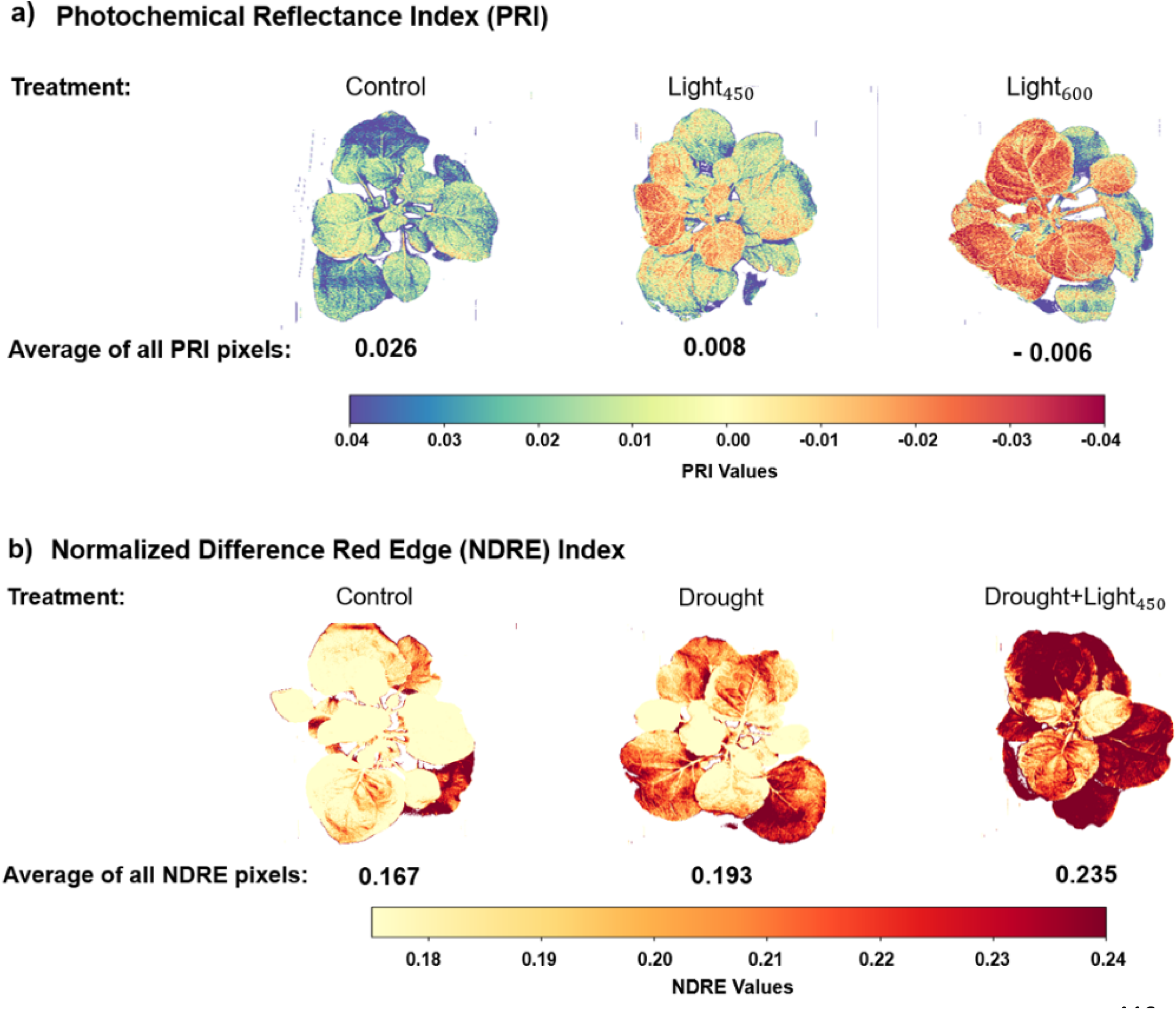
Responses of six different plants to various abiotic treatments are visualized through PRI (light-use efficiency) and NDRE (chlorophyll content) indices. Panel (a) shows PRI with a blue (high PRI) to red (low PRI) gradient. The increase in redness from Control to Light600 denotes a decrease in PRI as light stress increases. Panel (b) shows NDRE using a yellow (low NDRE) to red (high NDRE) gradient, where NDRE values are lowest in control plants, intermediate in Drought plants, and highest in Drought+Light450 plants.

For the three ML models, **Table 5** provides the prediction accuracies at different plant growth stages. The 3D+HSI-model outperformed both the HSI- and the 3D-models for stress identification at all growth stages. The improvement in model accuracy with plant age is consistent with the progressive accumulation of treatment-specific phenotypic signatures. At earlier stages, stress-induced differences in canopy structure, pigment status and photochemical performance were still developing, resulting in greater overlap among treatments. By 31–34 DAS, the effects of the imposed environmental treatments on traits such as PRI, NDRE, digital biomass and NPCI had become more pronounced, improving class separability (Figure 3).

**Table 5.** Accuracy of the 3D-, HSI- and 3D+HSI-models at various plant growth stages (DAS = days after sowing). Values in bold indicate the highest accuracy for identifying plants grown in a given stress condition at each time point.

| DAS | Overall test accuracy (%) |  |  |
| --- | --- | --- | --- |
|  | 3D-model | HSI-model | 3D + HSI-model |
| 27 | 62 | 64 | **78** |
| 27-29 | 70 | 69 | **85** |
| 27-31 | 71 | 73 | **86** |
| 27-34 | 72 | 81 | **89** |

An additional temporal-ablation analysis showed that the fused model using the complete 27–34 DAS feature set performed significantly better than models restricted to measurements from either 31 DAS or 34 DAS alone after Holm correction (Appendix 8). Thus, serial measurements provided predictive information beyond either late measurement alone. Because treatments continued throughout the scanning period, this result demonstrates the value of integrating phenotypic measurements collected across plant development under ongoing treatment exposure.

Overall, the 3D+HSI model showed strong classification performance, with low confusion among treatments (see Appendix 5) and high balanced accuracy across stress classes (see Table 4). Therefore, it is possible to classify single and combined stresses using the 3D+HSI traits.

### Test results with confidence scores for the model using combined traits (3D+HSI-model)

Figure 5 illustrates confidence for one correctly classified plant. Across all 108 out-of-fold predictions, correct classifications had substantially higher maximum confidence than incorrect classifications (median 0.953 versus 0.677; two-sided Mann–Whitney p < 0.001), indicating that model confidence generally tracked prediction correctness. Misclassifications occurred across all four experiments rather than being confined to one experiment. Detailed confidence distributions and error counts are provided in Appendi× 10.

**Figure 5.**
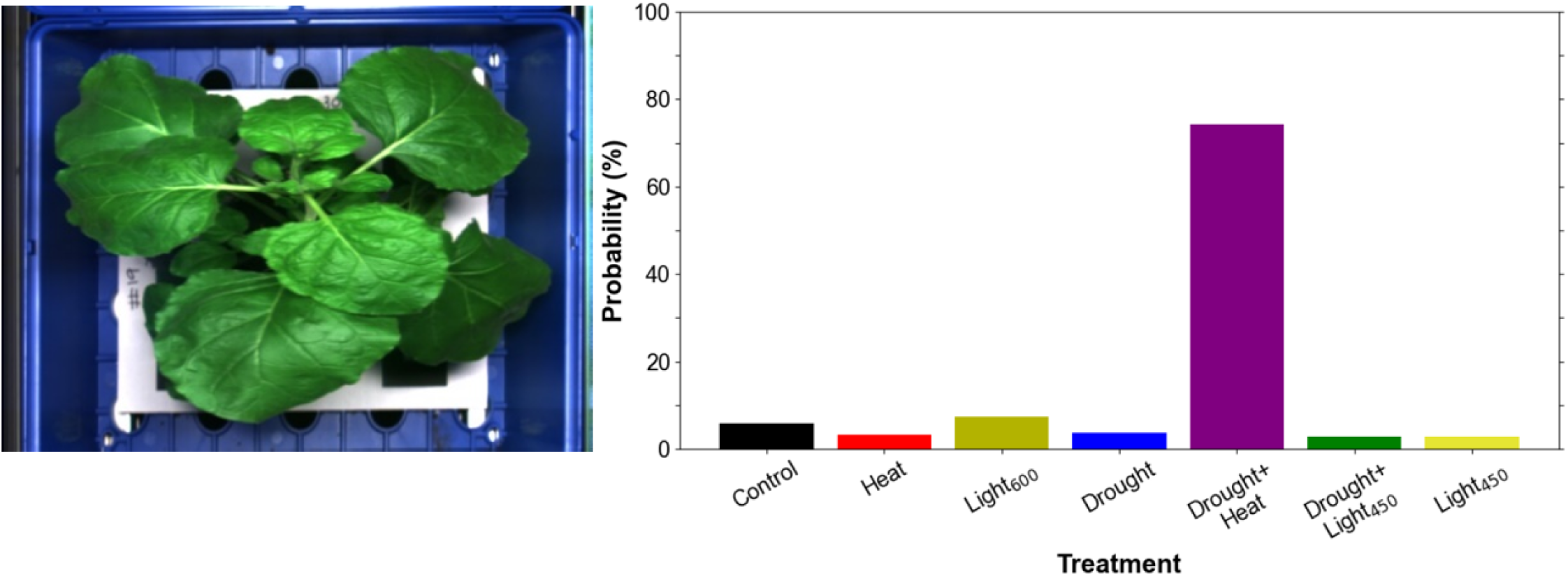
Predictions of the 3D + HSI ML model with probability scores at 34 days after sowing (DAS). The plant shown on the left was exposed to the Drought + Heat stress. The bar graph on the right depicts the probabilities, shown as percentages, of the plant to belong to one or the other abiotic treatments. The model assigned the highest confidence to Drought+Heat, which was the treatment imposed on this plant during growth.

## Discussion

The results of our study demonstrate that it is possible to identify the specific type of stress to which a plant had been exposed during growth, including distinguishing among multiple stressors which typically induce superficially similar changes in plant phenotype (e.g., high heat and drought), by comparing ML models developed using data from 2D-hyperspectral imaging and 3D-multispectral scans. The ability to identify the stresses experienced by plants may have value for commercial operations (e.g. horticulture and broadacre cropping) where understanding the drivers of variations in growth and chemical composition is important. However, further validation across independent experiments, species and growing environments will be required to assess the broader applicability of these approaches. In addition to stress events leaving a persistent signature at the molecular level (Lin, Chai et al. 2014, Lämke and Bäurle 2017), plants often exhibit a suite of anatomical, physiological and biochemical phenotypes in response to environmental conditions (Atkin, Loveys et al. 2005, Wright, Reich et al. 2006, Poorter, Niinemets et al. 2009, Nicotra, Atkin et al. 2010, Asao, Hayes et al. 2020, Wang, Townsend et al. 2022). Our study demonstrates that these phenotypic changes can be characterised via high-throughput phenotyping tools, and that the resulting data can be used to accurately classify plants according to their environmental conditions within the experimental system investigated here using a combination of high-throughput, off-the-shelf 2D hyperspectral and 3D-multispectral scanners. While past phenotyping studies have typically relied on either multispectral or hyperspectral techniques (Cheshkova 2022), our results highlight that combining data from these two tools, one providing deeper insights into shoot chemical composition and the other information on shoot structure and pigment composition, improves the identification of environmental treatments and provides complementary information on plant growth and performance compared with either technique alone.

### Phenotypic responses to environmental changes

Key to the training of the ML models was the creation of a wide range of morphological and chemical phenotypes in the shoot of *N. benthamiana* plants and the use of 2D-hyperspectral and 3D-multispectral scans during plant development. Changes in growth irradiance, water availability and temperature can alter the chemical composition of leaves in ways that ultimately affect their reflectance properties. For example, patterns of N allocation within leaves (including within the photosynthetic system) are altered by growth irradiance, along with shifts in concentrations of soluble phenolics (Poorter, Pepin et al. 2006). Drought stressed plants also exhibit significant changes in chemical composition, marked by changes in leaf water content, along with increases in compatible solute concentrations (e.g. proteins and amino acids, carbohydrates and organic acids) (Chaves, Maroco et al. 2003). Similarly, increases in growth temperature often result in reduced leaf N concentrations, underpinned by a reduction in the abundance of the carbon dioxide (CO_2_)-fixing enzyme, ribulose-1,5-bisphosphate carboxylase/oxygenase (RubisCO) (Scafaro, Xiang et al. 2017). Increasing growth temperatures can also alter the proportion of leaf N allocated to non-photosynthetic cell components (e.g. cell wall N and defence compounds) (Yamori, Noguchi et al. 2005), N partitioning between photosynthetic components (e.g. carboxylation capacity vs capacity for ribulose-1,5-bisphosphate (RuBP) regeneration (Hikosaka, Ishikawa et al. 2005)), and the abundance of soluble sugars and amino acids (Rashid, Scafaro et al. 2020). As an emergent property, these environmentally induced changes in leaf chemistry alter the absorbance and the reflectance of leaves, changing their overall multispectral and hyperspectral scan outputs (Figures 3 & 4, Table 4).

In addition to the changes in leaf chemistry, variations in shoot height, leaf surface and shoot biomass were induced when plants were grown under different irradiance, watering and temperature regimes (Figure 2). Growth under three different irradiances (300, 450 and 600 μmol photons m^−2^ s^−1^) did not affect plant height, but did significantly affect total leaf surface, while the drought treatment led to a substantial reduction in shoot biomass (Figure 2). Leaf area was also reduced by drought, particularly in heat-treated plants (Figure 2). While recent work has highlighted divergent responses of different *N. benthamiana* genotypes to drought (Asadyar, de Felippes et al. 2024), it seems likely that the mechanisms underpinning reduced growth under drought were reduced stomatal conductance (to limit water loss) with negative consequences for photosynthetic CO_2_ uptake. Growth under the heat treatment resulted in plants being taller, with more leaf area, but with no significant change in shoot biomass. Building on these observations, our study shows that an ML model trained on either 2D-HSI or 3D multispectral scanning data can distinguish between plants that were exposed to these different environments.

Past studies have shown that as part of a thermo-morphogenesis response (Casal and Balasubramanian 2019, Quint, Delker et al. 2023), the ratios of leaf area to leaf mass (Poorter, Niinemets et al. 2009) and of leaf area to stem elongation (Jensen, Eilertsen et al. 1996, Gray, Ostin et al. 1998, Casal and Balasubramanian 2019) increase with higher growth temperature, while heat can also lead to the upward bending of leaves (i.e. hyponastic movement) (Vile, Pervent et al. 2012, Sang, Fan et al. 2023, Thomas, Heckathorn et al. 2023). Such observations may account for the fact that heat-treated *N. benthamiana* plants were taller and had bigger leaf area (but not greater shoot mass) than plants grown under the control conditions. The fact that all three abiotic factors - irradiance, water supply and temperature - have an impact on canopy structure highlights the importance of the 3D scanning of shoots as part of a diagnostic approach to identify the specific type of stress, or combination of stresses, to which a plant has been exposed during growth.

### Model performance

Importantly, 2D-hyperspectral and 3D-multispectral imaging techniques applied in our study did not perform equally well for the different environmental treatments, likely due to differences in what they measure. For example, the 3D scanner provides estimates of morphological traits such as plant height and size, along with spectral indices such as NDVI. The 3D-model, trained on data from the 3D multispectral scanner, tended to place weight on information on plant height, digital biomass and spectral traits such as greenness (Table 4), and thus performed best on identifying plants from the Heat and the Drought+Heat treatments. These two stress treatments tended to affect plant height and biomass accumulation the most. By contrast, the HSI-model lacks 3D plant structural information, but collects considerably more information on leaf pigment concentrations and photosynthetic performance. The HSI-model outperformed the 3D-model by 20% to 30% when identifying plants submitted to the Drought treatment, the Light treatments (Light_450_ and Light_600_), and the combined Drought+Light_450_ treatment. This suggests that the lower performance of the 3D model for these treatments was not simply a modelling limitation but reflected the biological nature of the phenotypes: these treatments appeared to leave stronger spectral/biochemical signatures than whole-canopy structural signatures. Consistent with these observations, past studies have reported that drought, heat and altered irradiance can have pronounced impacts on chlorophyll and carotenoid ratios, pigment concentrations and photoprotective responses, even when effects on whole-canopy biomass accumulation are less pronounced (Lefsrud, Kopsell et al. 2006, Gallé and Feller 2007, Valladares and Niinemets 2008).

Combined-stress treatments are unlikely to produce phenotypes that are simply additive combinations of the corresponding single stresses. Drought, heat and high irradiance can interact through shared effects on stomatal conductance, leaf energy balance, pigment composition, photoprotection and carbon allocation (Rizhsky et al. 2004; Suzuki et al. 2014). For example, drought can limit transpirational cooling and carbon assimilation, while high light increases the need for photoprotective energy dissipation (Zhou et al. 2007; Suzuki et al. 2014). These interacting responses may explain why the Drought+Light450 and Drought+Heat classes were not always positioned between the corresponding single-stress classes and why combining structural and spectral information improved their identification.

While both the 2D-hyperspectral imaging and the 3D-multispectral scans could be used to differentiate environmental stress treatments imposed during growth, our study demonstrates that combining both 2D spectral and 3D morphological traits with ML provides a useful phenomics framework for more robustly characterising the complex range of modifications occurring at the leaf- and the canopy-level in response to stressful environmental conditions. Combining the modalities yielded the highest numerical within-study accuracy (88.9%). The fused model significantly outperformed the 3D-only model and showed directional evidence of improvement over the HSI-only model, although the latter comparison was not significant after two-sided Holm correction. This pattern is consistent with complementarity between structural and spectral information. This improvement was biologically plausible because the two modalities captured different components of the treatment-associated phenotype. The 3D-multispectral traits captured accumulated changes in canopy architecture, including plant height, projected area and digital biomass, whereas the HSI indices captured finer-scale variation in pigment status, photochemical performance and senescence-related spectral responses (Gamon et al. 1997; Garbulsky et al. 2011; Boiarskii and Hasegawa 2019; Merzlyak et al. 1999). The fused model therefore had access to both structural and biochemical/photochemical treatment signatures of the imposed environments. By leveraging the unique capabilities of each technique, the combined model could account for both structural traits and fine-scale changes in leaf biochemistry in response to the treatments, capturing an overall fingerprint of how plant traits vary across abiotic conditions.

The greater weighting assigned to PRI and NDRE is biologically consistent with the effects of the imposed stress treatments on leaf photochemistry and pigment status. PRI is commonly associated with photosynthetic light-use efficiency and changes in xanthophyll-cycle activity, and is therefore sensitive to irradiance and stress-induced changes in energy dissipation (Gamon et al. 1997; Garbulsky et al. 2011). NDRE is a red-edge index linked to chlorophyll status and canopy nitrogen/chlorophyll-related variation, making it useful for distinguishing treatments that alter pigment accumulation without necessarily producing strong structural differences (Boiarskii and Hasegawa 2019). PSRI and NPCI provided additional information on senescence-related pigment balance and chlorophyll/carotenoid relationships (Peñuelas et al. 1995; Merzlyak et al. 1999). Digital biomass complemented these spectral traits by capturing the accumulated structural consequences of the treatments. Thus, the fused model likely improved classification because it combined biochemical/photochemical stress signatures from HSI with structural treatment responses captured by the 3D scanner.

When comparing the parameters that were most heavily weighted in the models to differentiate plants from different treatments, the combined 3D+HSI-model did not necessarily prioritize the same parameters as did the models using only one of the two datasets. While the combined 3D+HSI-model most heavily weighted the same hyperspectral parameters as the HSI-model (PRI and NDRE), PSRI received the greatest weighting among the multispectral traits and was the fourth most weighted parameter in the HSI model. PSRI is calculated based on three wavelengths covering bands in the red, green and NIR part of the spectrum, while PRI and NDRE are based on two wavelengths each. It is therefore likely that PSRI carries complementary information to the other two indices. While the 3D-model had two metrics of plant size in its top three most weighted parameters (digital biomass and plant height), the combined 3D+HSI-model placed weight on one plant size metric (digital biomass) in the top eight weighted parameters. Thus, when the model had both structural and spectral data available to draw on, the spectral data provided greater insight into which treatments plants experienced.

One question that arises is whether it is necessary to use a hyperspectral sensor to gain information about how changes in the environment affect leaf chemical composition. Both the 2D- and 3D-datasets included spectral data that could provide insights into leaf physiological and biochemical performance (Behmann, Steinrücken et al. 2014, Silva-Perez, Molero et al. 2018, Coast, Shah et al. 2019), with the 3D multispectral scanner also providing information on the canopy structure. Given this, one possibility was that the 3D multispectral scanning model would perform better than the hyperspectral imaging model – however, this was not the case. Rather, the 3D multispectral imaging model only outperformed the hyperspectral imaging model for plants grown under the Heat and Drought+Heat treatments (Table 4), where plant height (and leaf area for the Heat treatment) were significantly enhanced compared to plants from other treatments. For all other treatments, HSI-models outperformed 3D-models (Table 4). Thus, what the HSI-model lacked in structural information, it more than made up for by leveraging a much richer spectral dataset than was available to the multispectral imaging model.

### Concluding comments

Looking forward, we see several challenges and opportunities. In our study, models were evaluated using plant-level leave-one-out testing within a controlled-environment dataset from one species. Although this approach maximised the use of the available data and avoided plant-level data leakage, it does not provide an external validation test across independent experiments, cultivars, species or production environments. Therefore, the reported accuracy should be interpreted as evidence for within-study classification performance rather than as a guarantee of direct field or commercial deployment. The present study used feature-level fusion of vegetation indices and 3D phenotypic parameters because this strategy is interpretable and reduces overfitting risk in a relatively small dataset. Future studies with larger datasets should revisit the full continuous hyperspectral signal using wavelength-importance analysis, PCA/PLS-based feature extraction and spectral feature-learning approaches. With sufficient sample sizes, CNNs, Transformers or other spectral deep-learning methods could be used to learn directly from raw spectra rather than relying only on predefined vegetation indices. More advanced multimodal fusion strategies, including early fusion, late fusion and attention-based fusion, could also be explored to better exploit complementarity between 3D structural and spectral information. Deeper insights into how environmental stresses affect plant structure and performance through time could likely be obtained by mapping 2D-hyperspectral images onto the 3D surface of plant canopies. Doing so would enable predictions to be made about where in the canopy an environmental stress is having the greatest effect, with that information potentially guiding how to manage and intervene to modulate the impacts of environmental stress. A further challenge is the need to increase the spatial and temporal scales over which sensors monitor canopies, noting that for industry, it will be important that there is an option to deploy non-invasive scanning tools across large-scale commercial glasshouses and field-production systems using sensors mounted on gantries or drones (with foundational datasets being needed to train ML models for each cropping system). In both cases, the challenges created by different light sources (diffuse and direct), humidity and occluded canopies will need to be addressed – all areas worthy of future investigation.

## Acknowledgements

The plant research infrastructure and phenotyping expertise was provided by the Australian Plant Phenomics Network at the Australian National University. The Australian Plant Phenomics Network is supported by the National Collaborative Research Infrastructure Strategy of the Australian Government. We acknowledge support from the ANU Centre for Entrepreneurial Agri-Technology (now ANU Agrifood Innovation Institute) and funding provided by Medicago Inc. (Québec, Canada). We also thank Prof Bob Furbank for comments on an earlier draft of the manuscript.

## CRediT authorship contribution statement

Frederike Stock: Methodology, Investigation, Formal analysis, Writing – original draft, Writing – review & editing.

Saswat Panda: Methodology, Data curation, Formal analysis, Visualization, Software, Validation, Writing – original draft, Writing – review & editing.

Richard Poiré: Supervision, Methodology, Resources, Writing – original draft, Writing – review & editing.

Timothy Brown: Supervision, Resources, Writing – original draft, Writing – review & editing.

Ayesha Akram: Investigation, Writing – review & editing.

Liang Zheng: Supervision, Methodology, Writing – review & editing.

Huan Lei: Methodology, Writing – review & editing.

Ruyi Zha: Data curation, Writing – review & editing.

Mingrui Zhao: Data curation, Writing – review & editing.

Sebastien Isabelle: Resources, Writing – review & editing.

Michèle Martel: Resources, Writing – review & editing.

Marc-André Comeau: Resources, Writing – review & editing.

Louis-Philippe Hamel: Resources, Writing – review & editing.

Pierre-Olivier Lavoie: Resources, Writing – review & editing.

Marc André D’Aoust: Resources, Funding acquisition, Writing – review & editing.

Hannah Reithinger: Resources, Writing – review & editing.

Pooja Saxena: Resources, Writing – review & editing.

Eric A. Stone: Formal analysis, Writing – review & editing.

Hongdong Li: Conceptualization, Supervision, Writing – review & editing.

Danielle A. Way: Supervision, Writing – original draft, Writing – review & editing.

Owen K. Atkin: Conceptualization, Supervision, Funding acquisition, Writing – original draft, Writing – review & editing.

## Data Availability

The processed feature datasets, model-training code and instructions required to reproduce the plant-level leave-one-out evaluation are available from the corresponding authors upon reasonable request. For peer-review purposes, a separate ZIP file was provided to the journal containing the processed feature dataset, analysis code, and instructions for reproducing the leave-one-out evaluation reported in this study.

## References

Afzal, A., S. W. Duiker and J. E. Watson (2017). “Leaf thickness to predict plant water status.” Biosystems Engineering 156: 148–156.

Asadyar, L., F. F. de Felippes, J. Bally, C. J. Blackman, J. An, F. C. Sussmilch, L. Moghaddam, B. Williams, S. J. Blanksby, T. J. Brodribb and P. M. Waterhouse (2024). “Evidence for within-species transition between drought response strategies in Nicotiana benthamiana.” New Phytologist 244(2): 464–476.

Asao, S., L. Hayes, M. J. Aspinwall, P. D. Rymer, C. Blackman, C. J. Bryant, D. Cullerne, J. J. G. Egerton, Y. Fan, P. Innes, A. H. Millar, J. Tucker, S. Shah, I. J. Wright, G. Yvon-Durocher, D. Tissue and O. K. Atkin (2020). “Leaf trait variation is similar among genotypes of Eucalyptus camaldulensis from differing climates and arises in plastic responses to the seasons rather than water availability.” New Phytologist 227(3): 780–793.

Atkin, O. K., B. R. Loveys, L. J. Atkinson and T. L. Pons (2005). “Phenotypic plasticity and growth temperature: understanding interspecific variability.” Journal of Experimental Botany 57(2): 267–281.

Behmann, J., J. Steinrücken and L. Plümer (2014). “Detection of early plant stress responses in hyperspectral images.” ISPRS Journal of Photogrammetry and Remote Sensing 93: 98–111.

Benjamini, Y. and Y. Hochberg (1995). “Controlling the false discovery rate: a practical and powerful approach to multiple testing.” Journal of the Royal Statistical Society: Series B (Methodological) 57(1): 289–300.

Boiarskii, B. and H. Hasegawa (2019). “Comparison of NDVI and NDRE indices to detect differences in vegetation and chlorophyll content.” Journal of Mechanics of Continua and Mathematical Sciences spl1.

Buckley, T. N. (2019). “How do stomata respond to water status?” New Phytologist 224(1): 21–36.

Cammarano, D., G. Fitzgerald, B. Basso, G. O’Leary, D. Chen, P. Grace and C. Fiorentino (2011). “Use of the Canopy Chlorophyl Content Index (CCCI) for remote estimation of wheat nitrogen content in rainfed environments.” Agronomy Journal 103(6): 1597–1603.

Casal, J. J. and S. Balasubramanian (2019). “Thermomorphogenesis.” Annual Review of Plant Biology 70: 321–346.

Cawley, G. C. and N. L. C. Talbot (2010). “On over-fitting in model selection and subsequent selection bias in performance evaluation.” Journal of Machine Learning Research 11: 2079–2107.

Charng, Y.-y., S. Mitra and S.-J. Yu (2022). “Maintenance of abiotic stress memory in plants: Lessons learned from heat acclimation.” The Plant Cell 35(1): 187–200.

Chaves, M. M., J. P. Maroco and J. S. Pereira (2003). “Understanding plant responses to drought - from genes to the whole plant.” Functional Plant Biology 30(3): 239–264.

Chen, T. and C. Guestrin (2016). XGBoost: A scalable tree boosting system. Proceedings of the 22nd ACM SIGKDD International Conference on Knowledge Discovery and Data Mining. San Francisco, California, USA, Association for Computing Machinery: 785–794.

Cheshkova, A. F. (2022). “A review of hyperspectral image analysis techniques for plant disease detection and identification.” Vavilovskii Zhurnal Genet Selektsii 26(2): 202–213.

Coast, O., S. Shah, A. Ivakov, O. Gaju, P. B. Wilson, B. C. Posch, C. J. Bryant, A. C. A. Negrini, J. R. Evans, A. G. Condon, V. Silva-Pérez, M. P. Reynolds, B. J. Pogson, A. H. Millar, R. T. Furbank and O. K. Atkin (2019). “Predicting dark respiration rates of wheat leaves from hyperspectral reflectance.” Plant, Cell & Environment 42(7): 2133–2150.

Cochran, W. G. (1950). “The comparison of percentages in matched samples.” Biometrika 37(3/4): 256–266.

Demmig-Adams, B. and W. W. Adams III (1992). “Carotenoid composition in sun and shade leaves of plants with different life forms.” Plant, Cell & Environment 15(4): 411–419.

Efron, B. (1979). “Bootstrap methods: another look at the jackknife.” The Annals of Statistics 7(1): 1–26.

Evans, J. R. and H. Poorter (2001). “Photosynthetic acclimation of plants to growth irradiance: the relative importance of specific leaf area and nitrogen partitioning in maximizing carbon gain.” Plant, Cell & Environment 24(8): 755–767.

Everingham, S. E., C. A. Offord, M. E. B. Sabot and A. T. Moles (2024). “Leaf morphological traits show greater responses to changes in climate than leaf physiological traits and gas exchange variables.” Ecology and Evolution 14(3): e10941.

Faqeerzada, M. A., E. Park, T. Kim, M. S. Kim, I. Baek, R. Joshi, J. Kim and B.-K. Cho (2023). “Fluorescence hyperspectral imaging for early diagnosis of heat-stressed ginseng plants.” Applied Sciences 13(1): 31.

Francini, A. and L. Sebastiani (2019). “Abiotic stress effects on performance of horticultural crops.” Horticulturae 5: 67.

Galieni, A., N. D’Ascenzo, F. Stagnari, G. Pagnani, Q. Xie and M. Pisante (2021). “Past and future of plant stress detection: An overview from remote sensing to positron emission tomography.” Frontiers in Plant Science 11.

Gallé, A. and U. Feller (2007). “Changes of photosynthetic traits in beech saplings (Fagus sylvatica) under severe drought stress and during recovery.” Physiologia Plantarum 131(3): 412–421.

Gamon, J. A., L. Serrano and J. S. Surfus (1997). “The photochemical reflectance index: an optical indicator of photosynthetic radiation use efficiency across species, functional types, and nutrient levels.” Oecologia 112(4): 492–501.

Garbulsky, M. F., J. Peñuelas, J. Gamon, Y. Inoue and I. Filella (2011). “The photochemical reflectance index (PRI) and the remote sensing of leaf, canopy and ecosystem radiation use efficiencies: A review and meta-analysis.” Remote Sensing of Environment 115(2): 281–297.

Gill, T., S. K. Gill, D. K. Saini, Y. Chopra, J. P. de Koff and K. S. Sandhu (2022). “A comprehensive review of high throughput phenotyping and Machine Learning for plant stress phenotyping.” Phenomics 2(3): 156–183.

Gorsuch, P. A., S. Pandey and O. K. Atkin (2010). “Thermal de-acclimation: how permanent are leaf phenotypes when cold-acclimated plants experience warming?” Plant, Cell & Environment 33(7): 1124–1137.

Gray, W. M., A. Ostin, G. Sandberg, C. P. Romano and M. Estelle (1998). “High temperature promotes auxin-mediated hypocotyl elongation in Arabidopsis.” Proc Natl Acad Sci U S A 95(12): 7197–7202.

Green, J. M., H. Appel, E. M. Rehrig, J. Harnsomburana, J.-F. Chang, P. Balint-Kurti and C.-R. Shyu (2012). “PhenoPhyte: a flexible affordable method to quantify 2D phenotypes from imagery.” Plant Methods 8(1): 45.

Grieve, B. D., T. Duckett, M. Collison, L. Boyd, J. West, H. Yin, F. Arvin and S. Pearson (2019). “The challenges posed by global broadacre crops in delivering smart agri-robotic solutions: A fundamental rethink is required.” Global Food Security 23: 116–124.

Hikosaka, K., K. Ishikawa, A. Borjigidai, O. Muller and Y. Onoda (2005). “Temperature acclimation of photosynthesis: mechanisms involved in the changes in temperature dependence of photosynthetic rate.” Journal of Experimental Botany 57(2): 291–302.

Holm, S. (1979). “A simple sequentially rejective multiple test procedure.” Scandinavian Journal of Statistics 6(2): 65–70.

Hoshino, R., Y. Yoshida and H. Tsukaya (2019). “Multiple steps of leaf thickening during sun-leaf formation in Arabidopsis.” The Plant Journal 100(4): 738–753.

Hughes, G. F. (1968). “On the mean accuracy of statistical pattern recognizers.” IEEE Transactions on Information Theory 14(1): 55–63.

Jensen, E., S. Eilertsen, A. Ernsten, O. Juntilla and R. Moe (1996). “Thermoperiodic control of stem elongation and endogenous gibberellins in Campanula isophylla.” Journal of Plant Growth Regulation 15(4): 167–171.

Lambers, H. and R. S. Oliveira (2019). Biotic influences: Interactions among plants. Plant Physiological Ecology. H. Lambers and R. S. Oliveira. Cham, Springer International Publishing: 615–648.

Lämke, J. and I. Bäurle (2017). “Epigenetic and chromatin-based mechanisms in environmental stress adaptation and stress memory in plants.” Genome Biology 18(1): 124.

Lazarević, B., K. Carović-Stanko, T. Safner and M. Poljak (2022). “Study of high-temperature-induced morphological and physiological changes in potato using nondestructive plant phenotyping.” Plants 11(24): 3534.

Lefsrud, M. G., D. A. Kopsell, D. E. Kopsell and J. Curran-Celentano (2006). “Irradiance levels affect growth parameters and carotenoid pigments in kale and spinach grown in a controlled environment.” Physiologia Plantarum 127(4): 624–631.

Li, Z., R. Guo, M. Li, Y. Chen and G. Li (2020). “A review of computer vision technologies for plant phenotyping.” Computers and Electronics in Agriculture 176: 105672.

Lin, M.-y., K.-h. Chai, S.-s. Ko, L.-y. Kuang, H.-S. Lur and Y.-y. Charng (2014). “A positive feedback loop between HEAT SHOCK PROTEIN101 and HEAT STRESS-ASSOCIATED 32-KD PROTEIN modulates long-term acquired thermotolerance illustrating diverse heat stress responses in rice varieties “ Plant Physiology 164(4): 2045–2053.

Liu, H., B. Bruning, T. Garnett and B. Berger (2020). “Hyperspectral imaging and 3D technologies for plant phenotyping: From satellite to close-range sensing.” Computers and Electronics in Agriculture 175: 105621.

Lobell, D. B. and S. M. Gourdji (2012). “The influence of climate change on global crop productivity.” Plant Physiology 160(4): 1686–1697.

Lowe, A., N. Harrison and A. P. French (2017). “Hyperspectral image analysis techniques for the detection and classification of the early onset of plant disease and stress.” Plant Methods 13(1): 80.

Lu, R. and Y.-R. Chen (1999). “Hyperspectral imaging for safety inspection of food and agricultural products.” Proc. SPIE 3544, Pathogen Detection and Remediation for Safe Eating 3544.

Mann, H. B. and D. R. Whitney (1947). “On a test of whether one of two random variables is stochastically larger than the other.” The Annals of Mathematical Statistics 18(1): 50–60.

MathWorks documentation. “Color-Based Segmentation Using the L*a*b* Color Space.” Retrieved 22/09/2023, 2023, from https://www.mathworks.com/help/images/color-based-segmentation-using-the-l-a-b-color-space.html#.

Melandri, G., K. R. Thorp, C. Broeckling, A. L. Thompson, L. Hinze and D. Pauli (2021). “Assessing drought and heat stress-induced changes in the cotton leaf metabolome and their relationship with hyperspectral reflectance.” Frontiers in Plant Science 12.

McNemar, Q. (1947). “Note on the sampling error of the difference between correlated proportions or percentages.” Psychometrika 12(2): 153–157.

Merzlyak, M. N., A. A. Gitelson, O. B. Chivkunova and V. Y. Rakitin (1999). “Non-destructive optical detection of pigment changes during leaf senescence and fruit ripening.” Physiologia Plantarum 106(1): 135–141.

Mohanty, S. P., D. P. Hughes and M. Salathé (2016). “Using Deep Learning for Image-Based Plant Disease Detection.” Frontiers in Plant Science 7.

Mukarram, M., S. Choudhary, D. Kurjak, A. Petek and M. M. A. Khan (2021). “Drought: Sensing, signalling, effects and tolerance in higher plants.” Physiologia Plantarum 172(2): 1291–1300.

Nicotra, A. B., O. K. Atkin, S. P. Bonser, A. M. Davidson, E. J. Finnegan, U. Mathesius, P. Poot, M. D. Purugganan, C. L. Richards, F. Valladares and M. van Kleunen (2010). “Plant phenotypic plasticity in a changing climate.” Trends in Plant Science 15(12): 684–692.

Niculescu-Mizil, A. and R. Caruana (2005). Predicting good probabilities with supervised learning. Proceedings of the 22nd International Conference on Machine Learning. Bonn, Germany, Association for Computing Machinery: 625–632.

Nobel, P. S. and S. P. Long (1985). Chapter 4 - Canopy Structure and Light Interception. Techniques in bioproductivity and photosynthesis (Second Edition). J. Coombs, D. O. Hall, S. P. Long and J. M. O. Scurlock, Pergamon: 41–49.

Ordonez, J., P. Bodegom, J.-P. Witte, I. Wright, P. Reich and R. Aerts (2009). “A global study of relationships between leaf traits, climate and soil measures of nutrient fertility.” Global Ecology and Biogeography 18: 137–149.

Otsu, N. (1979). “A threshold selection method from gray-level histograms.” IEEE Transactions on Systems, Man, and Cybernetics 9(1): 62–66.

Pedregosa, F., G. Varoquaux, A. Gramfort, V. Michel, B. Thirion, O. Grisel, M. Blondel, P. Prettenhofer, R. Weiss, V. Dubourg, J. Vanderplas, A. Passos, D. Cournapeau, M. Brucher, M. Perrot and E. Duchesnay (2011). “Scikit-learn: Machine Learning in Python.” Journal of Machine Learning Research 12: 2825–2830.

Peñuelas, J., F. Baret and I. Filella (1995). “Semi-empirical indices to assess carotenoids/chlorophyll-a ratio from leaf spectral reflectance.” Photosynthetica 31(2): 221–230.

Phenospex. “PlantEye F600 - Multispectral 3D scanner for plant phenotyping.” Retrieved 28/11/2023, 2023, from https://phenospex.com/products/plant-phenotyping/planteye-f600-multispectral-3d-scanner-for-plants/.

Poorter, H., Ü. Niinemets, L. Poorter, I. J. Wright and R. Villar (2009). “Causes and consequences of variation in leaf mass per area (LMA): a meta-analysis.” New Phytologist 182(3): 565–588.

Poorter, H., S. Pepin, T. Rijkers, Y. de Jong, J. R. Evans and C. Körner (2006). “Construction costs, chemical composition and payback time of high- and low-irradiance leaves.” Journal of Experimental Botany 57(2): 355–371.

Quint, M., C. Delker, S. Balasubramanian, M. Balcerowicz, J. J. Casal, C. D. M. Castroverde, M. Chen, X. Chen, I. De Smet, C. Fankhauser, K. A. Franklin, K. J. Halliday, S. Hayes, D. Jiang, J. H. Jung, E. Kaiserli, S. V. Kumar, D. Maag, E. Oh, C. M. Park, S. Penfield, G. Perrella, S. Prat, R. S. Reis, P. A. Wigge, B. C. Willige and M. van Zanten (2023). “25 Years of thermomorphogenesis research: milestones and perspectives.” Trends in Plant Science 28(10): 1098–1100.

Rashid, F. A. A., A. P. Scafaro, S. Asao, R. Fenske, R. C. Dewar, J. Masle, N. L. Taylor and O. K. Atkin (2020). “Diel- and temperature-driven variation of leaf dark respiration rates and metabolite levels in rice.” New Phytologist 228(1): 56–69.

Rizhsky, L., H. Liang, J. Shuman, V. Shulaev, S. Davletova and R. Mittler (2004). “When defense pathways collide. The response of Arabidopsis to a combination of drought and heat stress.” Plant Physiology 134(4): 1683–1696.

Sang, Q., L. Fan, T. Liu, Y. Qiu, J. Du, B. Mo, M. Chen and X. Chen (2023). “MicroRNA156 conditions auxin sensitivity to enable growth plasticity in response to environmental changes in Arabidopsis.” Nature Communications 14(1): 1449.

Scafaro, A. P., S. Xiang, B. M. Long, N. H. A. Bahar, L. K. Weerasinghe, D. Creek, J. R. Evans, P. B. Reich and O. K. Atkin (2017). “Strong thermal acclimation of photosynthesis in tropical and temperate wet-forest tree species: the importance of altered Rubisco content.” Global Change Biology 23(7): 2783–2800.

Schindelin, J., I. Arganda-Carreras, E. Frise, V. Kaynig, M. Longair, T. Pietzsch, S. Preibisch, C. Rueden, S. Saalfeld, B. Schmid, J.-Y. Tinevez, D. J. White, V. Hartenstein, K. Eliceiri, P. Tomancak and A. Cardona (2012). “Fiji: an open-source platform for biological-image analysis.” Nature Methods 9(7): 676–682.

Shahinfar, S., P. Meek and G. Falzon (2020). ““How many images do I need?” Understanding how sample size per class affects deep learning model performance metrics for balanced designs in autonomous wildlife monitoring.” Ecological Informatics 57: 101085.

Sheikh, M., F. Iqra, H. Ambreen, K. A. Pravin, M. Ikra and Y. S. Chung (2024). “Integrating artificial intelligence and high-throughput phenotyping for crop improvement.” Journal of Integrative Agriculture 23(6): 1787–1802.

Silva-Perez, V., G. Molero, S. P. Serbin, A. G. Condon, M. P. Reynolds, R. T. Furbank and J. R. Evans (2018). “Hyperspectral reflectance as a tool to measure biochemical and physiological traits in wheat.” J Exp Bot 69(3): 483–496.

Suzuki, N., R. M. Rivero, V. Shulaev, E. Blumwald and R. Mittler (2014). “Abiotic and biotic stress combinations.” New Phytologist 203(1): 32–43.

Terashima, I., Y. T. Hanba, Y. Tazoe, P. Vyas and S. Yano (2005). “Irradiance and phenotype: comparative eco-development of sun and shade leaves in relation to photosynthetic CO2 diffusion.” Journal of Experimental Botany 57(2): 343–354.

Thomas, M. D., S. A. Heckathorn and J. K. Boldt (2023). “Elevated CO2 Increases Severity of Thermal Hyponasty in Leaves of Tomato.” Horticulturae 9(8): 907.

Valladares, F. and Ü. Niinemets (2008). “Shade Tolerance, a Key Plant Feature of Complex Nature and Consequences.” Annual Review of Ecology Evolution and Systematics 39: 237–257.

Varma, S. and R. Simon (2006). “Bias in error estimation when using cross-validation for model selection.” BMC Bioinformatics 7: 91.

Vasseur, F., M. A.-O. Exposito-Alonso, O. J. Ayala-Garay, G. Wang, B. J. Enquist, D. Vile, C. Violle and D. A.-O. Weigel (2018). “Adaptive diversification of growth allometry in the plant Arabidopsis thaliana.” Proc Natl Acad Sci U S A 115(13): 3416–3421.

Vergara-Diaz, O., T. Vatter, S. C. Kefauver, T. Obata, A. R. Fernie and J. L. Araus (2020). “Assessing durum wheat ear and leaf metabolomes in the field through hyperspectral data.” The Plant Journal 102(3): 615–630.

Vile, D., M. Pervent, M. Belluau, F. Vasseur, J. Bresson, B. Muller, C. Granier and T. Simonneau (2012). “Arabidopsis growth under prolonged high temperature and water deficit: independent or interactive effects?” Plant Cell Environ 35(4): 702–718.

Virtanen, P., R. Gommers, T. E. Oliphant, M. Haberland, T. Reddy, D. Cournapeau, E. Burovski, P. Peterson, W. Weckesser, J. Bright, S. J. van der Walt, M. Brett, J. Wilson, K. J. Millman, N. Mayorov, A. R. J. Nelson, E. Jones, R. Kern, E. Larson, C. J. Carey, İ. Polat, Y. Feng, E. W. Moore, J. VanderPlas, D. Laxalde, J. Perktold, R. Cimrman, I. Henriksen, E. A. Quintero, C. R. Harris, A. M. Archibald, A. H. Ribeiro, F. Pedregosa, P. van Mulbregt, A. Vijaykumar, A. P. Bardelli, A. Rothberg, A. Hilboll, A. Kloeckner, A. Scopatz, A. Lee, A. Rokem, C. N. Woods, C. Fulton, C. Masson, C. Häggström, C. Fitzgerald, D. A. Nicholson, D. R. Hagen, D. V. Pasechnik, E. Olivetti, E. Martin, E. Wieser, F. Silva, F. Lenders, F. Wilhelm, G. Young, G. A. Price, G.-L. Ingold, G. E. Allen, G. R. Lee, H. Audren, I. Probst, J. P. Dietrich, J. Silterra, J. T. Webber, J. Slavič, J. Nothman, J. Buchner, J. Kulick, J. L. Schönberger, J. V. de Miranda Cardoso, J. Reimer, J. Harrington, J. L. C. Rodríguez, J. Nunez-Iglesias, J. Kuczynski, K. Tritz, M. Thoma, M. Newville, M. Kümmerer, M. Bolingbroke, M. Tartre, M. Pak, N. J. Smith, N. Nowaczyk, N. Shebanov, O. Pavlyk, P. A. Brodtkorb, P. Lee, R. T. McGibbon, R. Feldbauer, S. Lewis, S. Tygier, S. Sievert, S. Vigna, S. Peterson, S. More, T. Pudlik, T. Oshima, T. J. Pingel, T. P. Robitaille, T. Spura, T. R. Jones, T. Cera, T. Leslie, T. Zito, T. Krauss, U. Upadhyay, Y. O. Halchenko, Y. Vázquez-Baeza and C. SciPy (2020). “SciPy 1.0: fundamental algorithms for scientific computing in Python.” Nature Methods 17(3): 261–272.

Wang, Z., P. A. Townsend and E. L. Kruger (2022). “Leaf spectroscopy reveals divergent inter- and intra-species foliar trait covariation and trait–environment relationships across NEON domains.” New Phytologist 235(3): 923–938.

Welch, B. L. (1947). “The generalization of ‘Student’s’ problem when several different population variances are involved.” Biometrika 34(1-2): 28–35.

Wong Man Sing, C., C. Tang Hon Wai, L. Lam Lik Shan, H.-h. Cheng, K.-y. Hung, C. Kwok Yin-tung and C. Tang Tin-hang (2020). Introduction to the applications of remote sensing techniques on the tree health monitoring, The Hong Kong Polytechnic University, Department of Land Surveying and Geo-Informatics.

Wright, I. J., P. B. Reich, O. K. Atkin, C. H. Lusk, M. G. Tjoelker and M. Westoby (2006). “Irradiance, temperature and rainfall influence leaf dark respiration in woody plants: evidence from comparisons across 20 sites.” New Phytologist 169(2): 309–319.

Yamori, W., K. Noguchi and I. Terashima (2005). “Temperature acclimation of photosynthesis in spinach leaves: analyses of photosynthetic components and temperature dependencies of photosynthetic partial reactions.” Plant, Cell & Environment 28(4): 536–547.

Zhou, Y., H. M. Lam and J. Zhang. “Inhibition of photosynthesis and energy dissipation induced by water and high light stresses in rice.” Journal of Experimental Botany 58(5): 1207–1217.

