## Appendix for "Combining 3D-multispectral and hyperspectral imaging to identify environmental stress treatments imposed during plant growth"

### Supplementary Information

#### Appendix 1. Spectral light composition in Experiments 2 and 3

Light curves for experiments 2 and 3 are presented below to indicate spectral differences between Control and light450 and Light600 conditions.

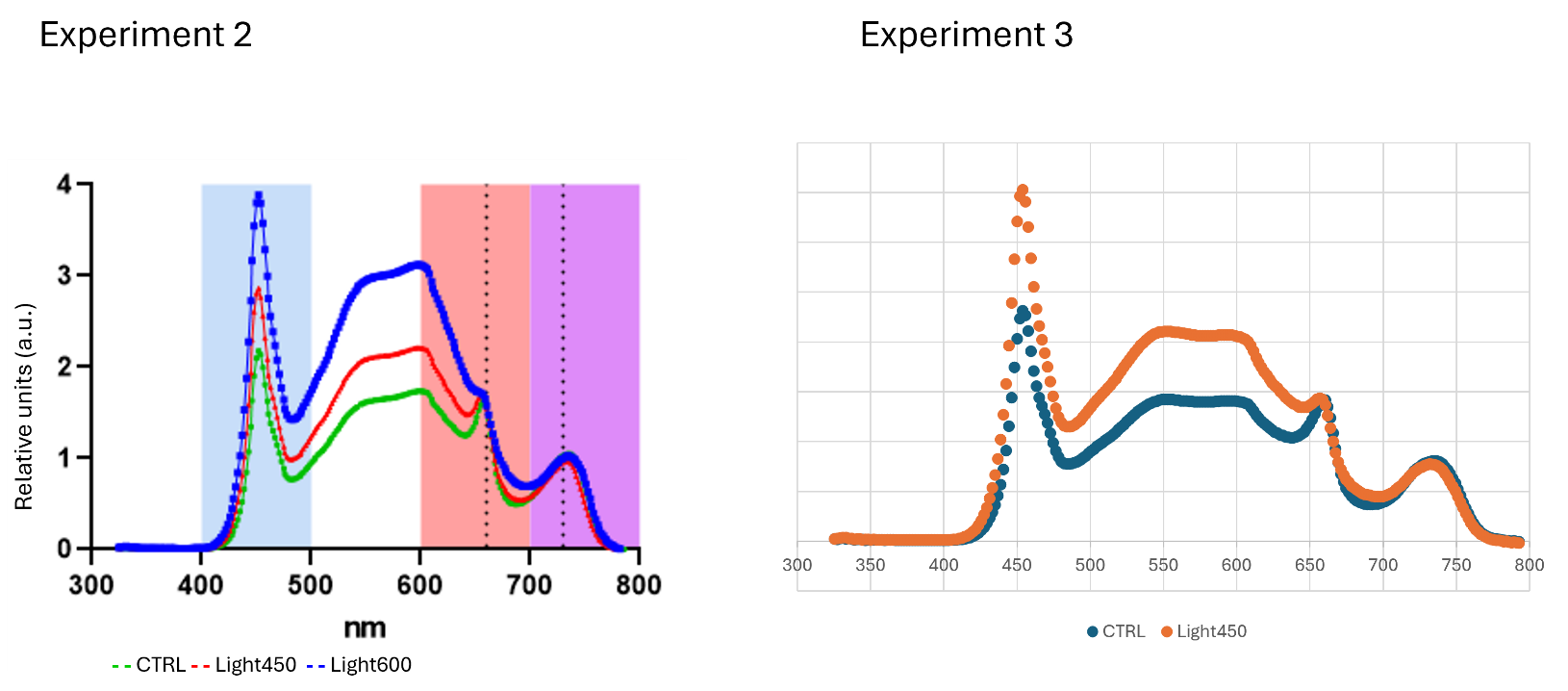

#### Appendix 2. Linear Interpolation

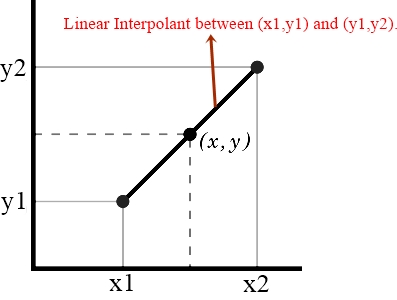

In order to create new data points within the range of a discrete set of existing data points, a curve fitting technique using linear polynomials is used, known as linear interpolation.

The formula used to calculate linear interpolation is below:

$$y\left( x \right) =y1+\left( x -x1 \right)\times\frac{(y2-y1)}{(x2-x1)}$$

For example, if value of PRI is missing on DAS 29, then we substitute above formula as below:

$$PRI (DAS 29) = PRI (DAS 27) + (29 -27) \times\frac{PRI(DAS 31) - PRI(DAS 27)}{(31-27)}$$

So, in our case, the 𝑥 in the formula represents time, and 𝑦 represents the trait value.

#### Appendix 3. Green colour detection and Otsu’s thresholding

To detect green colour, we first transformed the image from RGB colour space to CIE Lab colour space, which is a better representation of the human perception of colours and the preferred choice for identifying different colours in an image (MathWorks documentation). In the CIE Lab colour space, green colour is located in the A-channel, therefore we applied Otsu’s thresholding (an automatic thresholding method) (Otsu 1979) only to the A-channel to minimize small noises which are green but not the plant to obtain the plant mask.

#### Appendix 4. Leave one out testing

*Leave one out* testing is a special case of *leave-p-out* testing, which uses p samples as the test set and the remaining samples as the training set. On a test set of p samples and a training set, this is repeated for all possible ways to trim the original sample. The number of times the model gets trained is:

$∁(n,p)$ (n = total number of samples, p = number of samples chosen from the training set)

which becomes computationally very expensive, for p>1. Therefore, we used a special case of *leave-p-out* testing, known as *leave one out* testing. In *leave one out* testing p is 1, which means the model is trained and tested n times. The model performance is then evaluated by averaging the results from each iteration.

#### Appendix 5. Confusion matrices

Confusion matrices, List of traits with Model weightings for identification of single and combined stresses at DAS 34.

***3D Model (12 selected 3D-mutlispectral traits are used)***

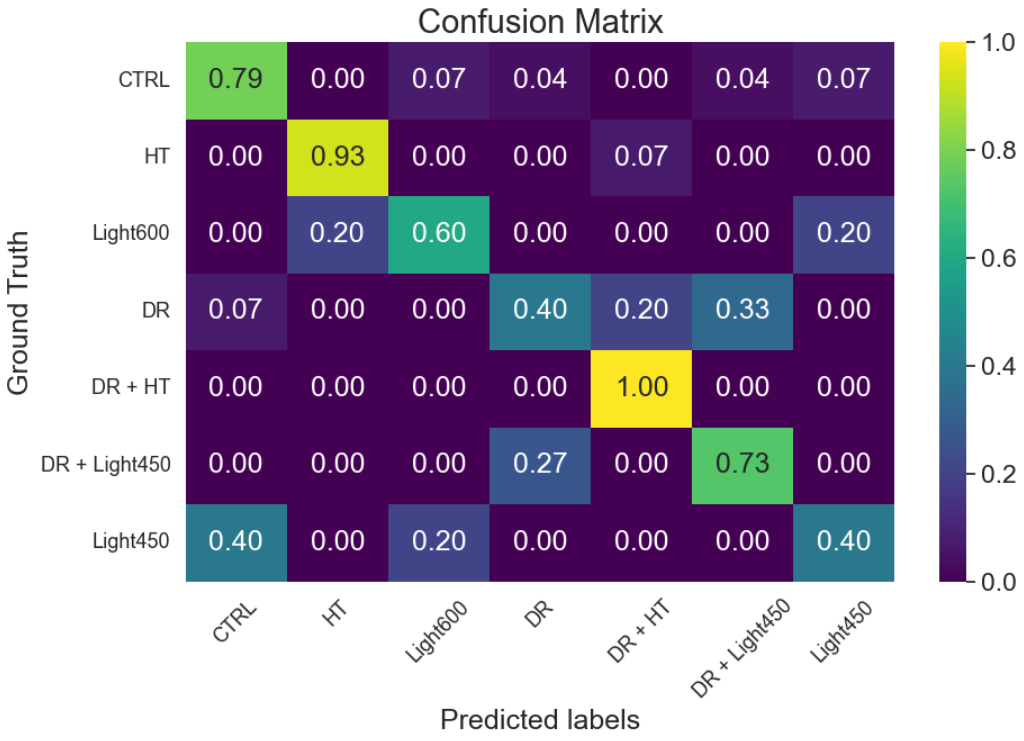

High confusion between Dr and Dr+Light_450_ was observed ((0.33+0.27)/2 = 0.3). Values are highlighted in red boxes in the confusion matrix.

**Feature Importance (Model weighting)**

| **Trait** | **Weighting (%)** |
| --- | --- |
| Digital biomass (mm³) | 13.82 |
| Greenness average | 12.38 |
| Height (mm) | 12.16 |
| PSRI average | 11.80 |
| Height Max (mm) | 10.98 |
| Leaf angle (°) | 8.36 |
| Leaf area (projected) (mm²) | 7.33 |
| Leaf area index (mm²/mm²) | 7.01 |
| Hue average (°) | 6.51 |
| NDVI average | 5.03 |
| Light penetration depth (mm) | 2.73 |
| NPCI average | 1.91 |

The table above lists all the traits used in the 3D model and their weightings.

***HSI Model (All 11 hyperspectral traits (indices) extracted were used)***

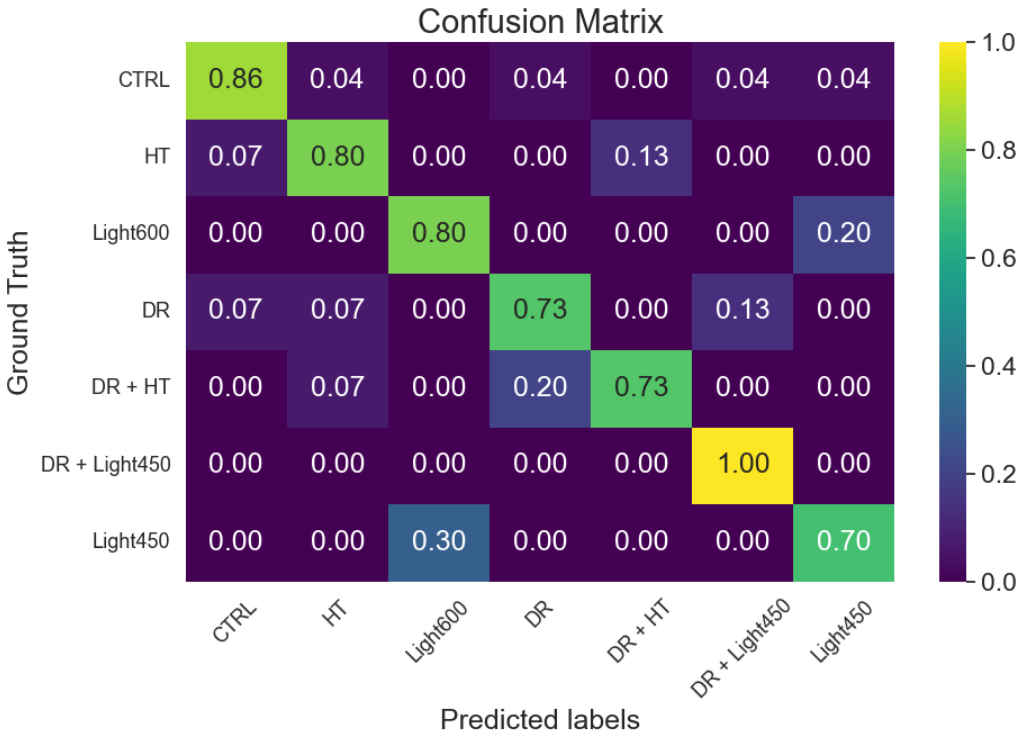

Confusion between Dr and Dr+Light_450_ was reduced ((0.13+0.00)/2 = 0.065). Values are highlighted in red boxes in the confusion matrix.

**Feature Importance (Model weighting)**

| **Trait** | **Weighting (%)** |
| --- | --- |
| PRI_avg | 20.53 |
| NDRE_avg | 19.84 |
| RNDVI_avg | 14.31 |
| NPCI_avg | 10.15 |
| NDVI_avg | 9.13 |
| CRI2_avg | 8.71 |
| NPQI_avg | 5.26 |
| PSRI_avg | 5.26 |
| SR_avg | 3.73 |
| SIPI_avg | 1.78 |
| CCCI_avg | 1.30 |

The table above lists all the traits used in the HSI model and their weightings.

***3D+HSI Model (Selected plant traits were used, 11*** 3D-multispectral ***traits and 4 HSI traits)***

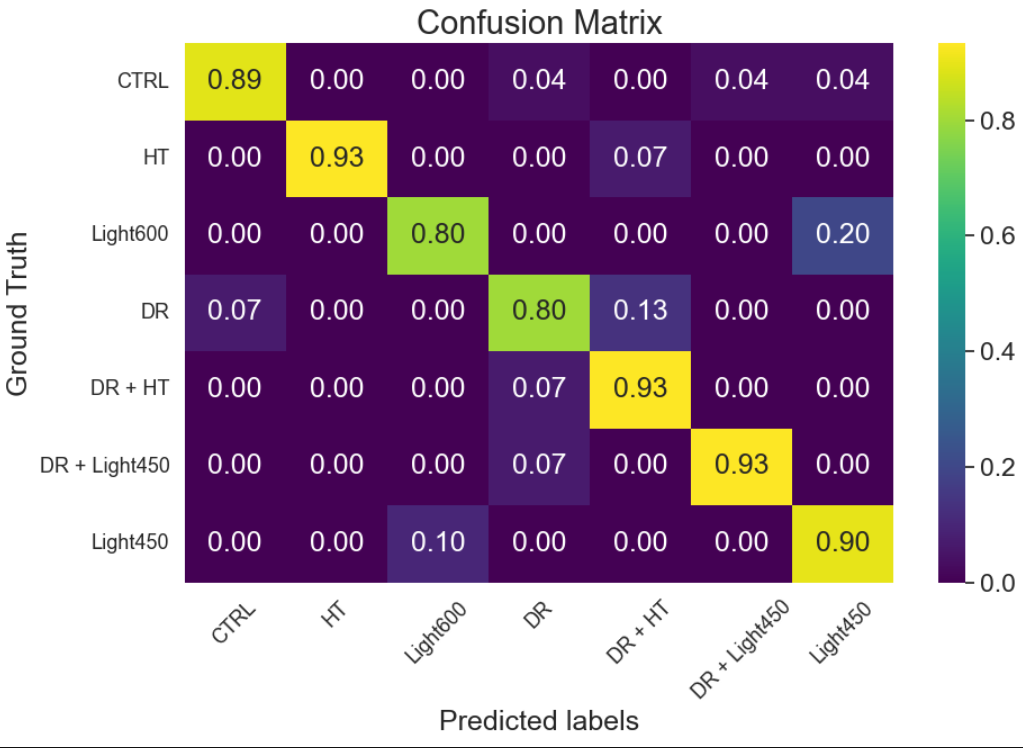

From the above confusion matrix, it was observed that the highest confusion was between Light_600_ and Light_450_ ((0.2+0.1)/2=0.15), highlighted in light blue boxes. In contrast, the confusion between Dr and Dr+Light_450_ was one of the lowest ((0.07+0.00)/2=0.035), highlighted in red boxes in the confusion matrix.

**Feature Importance (Model weighting)**

| **Source** | **Trait** | **Weighting (%)** |
| --- | --- | --- |
| HSI | PRI_avg | 15.57 |
| HSI | NDRE_avg | 15.20 |
| 3D | PSRI average | 14.84 |
| 3D | Digital biomass (mm³) | 11.29 |
| 3D | Greenness average | 7.43 |
| 3D | NPCI average | 7.20 |
| 3D | Leaf angle (°) | 6.89 |
| HSI | RENDVI_avg | 6.13 |
| 3D | Height (mm) | 3.95 |
| 3D | Leaf area (projected) (mm²) | 2.85 |
| 3D | NDVI average | 2.72 |
| 3D | Hue average (°) | 2.27 |
| HSI | NPCI_avg | 1.79 |
| 3D | Height Max (mm) | 0.98 |
| 3D | Leaf area index (mm²/mm²) | 0.88 |

The table above lists all the traits used in the 3D+HSI model and their weightings.

#### Appendix 6. Additional plant-level leave-one-out model-performance metrics and paired statistical comparisons

The tables below provide additional performance metrics for the three plant-level leave-one-out cross-validation models. Per-class precision, recall and F1-score are reported to account for class imbalance across stress treatments. Paired statistical tests were performed on the same 108 plant-level predictions to determine whether the observed differences among the three models were statistically supported.

Per-treatment precision, recall and F1-score for the plant-level leave-one-out cross-validation models.

| **Model** | **Treatment** | **Precision (%)** | **Recall (%)** | **F1-score (%)** |
| --- | --- | --- | --- | --- |
| 3D | CTRL | 81.5 | 78.6 | 80.0 |
| 3D | Heat | 87.5 | 93.3 | 90.3 |
| 3D | Light600 | 60.0 | 60.0 | 60.0 |
| 3D | Drought | 54.5 | 40.0 | 46.2 |
| 3D | Drought + Heat | 78.9 | 100.0 | 88.2 |
| 3D | Drought + Light450 | 64.7 | 73.3 | 68.8 |
| 3D | Light450 | 50.0 | 40.0 | 44.4 |
| HSI | CTRL | 92.3 | 85.7 | 88.9 |
| HSI | Heat | 80.0 | 80.0 | 80.0 |
| HSI | Light600 | 72.7 | 80.0 | 76.2 |
| HSI | Drought | 73.3 | 73.3 | 73.3 |
| HSI | Drought + Heat | 84.6 | 73.3 | 78.6 |
| HSI | Drought + Light450 | 83.3 | 100.0 | 90.9 |
| HSI | Light450 | 70.0 | 70.0 | 70.0 |
| 3D + HSI | CTRL | 96.2 | 89.3 | 92.6 |
| 3D + HSI | Heat | 100.0 | 93.3 | 96.6 |
| 3D + HSI | Light600 | 88.9 | 80.0 | 84.2 |
| 3D + HSI | Drought | 85.7 | 80.0 | 82.8 |
| 3D + HSI | Drought + Heat | 82.4 | 93.3 | 87.5 |
| 3D + HSI | Drought + Light450 | 93.3 | 93.3 | 93.3 |
| 3D + HSI | Light450 | 69.2 | 90.0 | 78.3 |

**Overall paired comparison of model correctness using Cochran's Q test.**

Cochran's Q test was used because the same 108 plants were evaluated by all three models, resulting in paired binary correctness outcomes. A significant result indicates that at least one model differs in classification performance (α=0.05).

| **Test** | **Statistic** | **df** | **p-value** |
| --- | --- | --- | --- |
| Cochran's Q | 11.619 | 2 | 0.002999 |

**Pairwise exact McNemar tests comparing paired model correctness.**

For each comparison, “A correct / B incorrect” counts plants correctly classified by the first model but incorrectly classified by the second model. “A incorrect / B correct” counts the reverse. Holm-adjusted p-values are reported for the two-sided tests; the one-sided directional p-value tests whether the first model in the comparison outperformed the second model (α=0.05).

| **Comparison** | **A correct / B incorrect** | **A incorrect / B correct** | **Discordant pairs** | **Two-sided exact McNemar p** | **Holm-adjusted two-sided p** | **One-sided directional p (A > B)** |
| --- | --- | --- | --- | --- | --- | --- |
| 3D+HSI vs 3D | 23 | 5 | 28 | 0.000912 | 0.002737 | 0.000456 |
| 3D+HSI vs HSI | 12 | 4 | 16 | 0.076813 | 0.153625 | 0.038406 |
| HSI vs 3D | 25 | 15 | 40 | 0.153860 | 0.153860 | 0.076930 |

**Bootstrap confidence intervals for paired accuracy differences.**

Bootstrap resampling was used to estimate uncertainty around paired accuracy differences. Confidence intervals that remain above zero provide directional support that the first model achieved higher accuracy than the second model.

| **Comparison** | **Observed accuracy difference** | **95% CI lower** | **95% CI upper** |
| --- | --- | --- | --- |
| 3D+HSI – 3D | 0.167 | 0.074 | 0.259 |
| 3D+HSI – HSI | 0.074 | 0.009 | 0.148 |
| HSI – 3D | 0.093 | -0.019 | 0.204 |

#### Appendix 7. Representative 75/25 held-out grid-search classifier comparison

This representative analysis used a 75/25 train/test split after grid-search tuning to compare XGBoost with Random Forest, Linear SVM and Logistic Regression. The table reports the best cross-validation macro F1-score and balanced accuracy from the training set, together with performance on the held-out test set of 27 plants. The hyperparameter search space used for the grid-search comparison is provided below the classifier-comparison table. This analysis was performed as a supplementary classifier-comparison and parameter-sensitivity analysis. It was not used to optimize or alter the model parameters used in the plant-level leave-one-out testing experiment reported in the main study. Because the held-out test set is small and depends on one representative split, the plant-level leave-one-out test analysis remains the primary performance estimate reported in the main manuscript.

**Representative classifier comparison using a 75/25 train/test split after grid-search tuning.**

| **Feature set** | **Classifier** | **Best CV macro F1-score (%)** | **Best CV balanced accuracy (%)** | **Held-out accuracy (%)** | **Held-out balanced accuracy (%)** | **Held-out macro F1-score (%)** |
| --- | --- | --- | --- | --- | --- | --- |
| 3D+HSI | XGBoost | 87.9 | 90.0 | 100.0 | 100.0 | 100.0 |
| 3D +HSI | Random Forest | 82.4 | 85.5 | 96.3 | 98.0 | 97.3 |
| 3D +HSI | Linear SVM | 85.5 | 87.6 | 96.3 | 98.0 | 97.3 |
| 3D +HSI | Logistic Regression | 88.8 | 90.5 | 96.3 | 98.0 | 97.3 |
| 3D | XGBoost | 59.5 | 62.3 | 74.1 | 70.9 | 70.3 |
| 3D | Random Forest | 65.7 | 70.4 | 70.4 | 63.8 | 62.7 |
| 3D | Linear SVM | 74.3 | 77.1 | 85.2 | 81.6 | 80.0 |
| 3D | Logistic Regression | 71.2 | 75.5 | 81.5 | 78.1 | 78.8 |
| HSI | XGBoost | 79.0 | 81.7 | 92.6 | 92.9 | 94.1 |
| HSI | Random Forest | 80.9 | 81.9 | 88.9 | 85.7 | 87.1 |
| HSI | Linear SVM | 83.1 | 84.0 | 81.5 | 73.0 | 71.0 |
| HSI | Logistic Regression | 84.0 | 84.9 | 77.8 | 74.5 | 74.4 |

**Hyperparameter search space used in the representative 75/25 grid-search classifier comparison.**

| **Classifier** | **Hyperparameter search space** |
| --- | --- |
| XGBoost | Two booster branches were searched: gblinear: booster = gblinear; n_estimators = [20, 30, 40, 50, 60]. gbtree: booster = gbtree; n_estimators = [100, 150, 200]; max_depth = [2, 3, 4]; learning_rate = [0.025, 0.05, 0.1]. |
| Random Forest | n_estimators = [100, 200, 500]; max_depth = [3, 4, 5, None]; min_samples_leaf = [1, 2, 4]; max_features = [sqrt, None]. |
| Linear SVM | Linear-kernel support vector classifier with standardisation. svc__C = [0.01, 0.1, 1.0, 10.0]. |
| Logistic Regression | Logistic regression with standardisation, solver = lbfgs, max_iter = 500. logisticregression__C = [0.01, 0.1, 1.0, 10.0]. |

#### Appendix 8. Temporal ablation of the fused model

Using the same fused-model configuration, temporal ablation compared all 60 trait-by-time features from 27–34 DAS with the 15 features from either 31 DAS or 34 DAS. The complete feature set performed significantly better than both single-date sets after Holm correction, and both paired-bootstrap 95% confidence intervals excluded zero (two-sided α = 0.05). This shows that serial measurements added predictive information beyond either late scan; because treatments remained in place throughout scanning, the analysis does not test post-treatment memory. Results are summarized in the tables below.

**Model-performance summary:**

| **Feature set** | **Features** | **Accuracy** | **Balanced accuracy** | **Macro F1** |
| --- | --- | --- | --- | --- |
| Complete 27–34 DAS History | 60 | 88.9% | 88.5% | 88.0% |
| 31 DAS only | 15 | 75.9% | 72.2% | 71.9% |
| 34 DAS only | 15 | 79.6% | 77.5% | 77.6% |

**Paired comparisons with the complete 27–34 DAS model:**

| **Comparison** | **History correct/single incorrect** | **History incorrect/single correct** | **Exact p** | **Holm p** | **Bootstrap 95% CI for difference** |
| --- | --- | --- | --- | --- | --- |
| History vs 31 DAS | 18 | 4 | 0.00434 | 0.00869 | 4.6 to 21.3 pp |
| History vs 34 DAS | 15 | 5 | 0.04139 | 0.04139 | 0.9 to 17.6 pp |

#### Appendix 9. Fold-wise batch-control-standardised

Sensitivity to batch- and chamber-associated control shifts was assessed by comparing the primary raw-feature LOOCV with fold-wise batch-control-standardised LOOCV, with normalization parameters estimated from training controls only. The accuracy difference was not significant according to Cochran's Q or the exact McNemar test, and the paired-bootstrap 95% confidence interval included zero (two-sided α = 0.05). These findings suggest that performance was not solely driven by batch-specific control differences, although residual batch and chamber confounding cannot be excluded. Results are summarized in the tables below.

**Model-performance summary:**

| **Feature treatment** | **Accuracy** | **Balanced accuracy** | **Macro F1** | **Plants** |
| --- | --- | --- | --- | --- |
| Raw features | 88.9% | 88.5% | 88.0% | 108 |
| Fold-wise batch-control standardized | 84.3% | 81.4% | 81.2% | 108 |

Paired statistical assessment:

| **Analysis** | **Statistic or paired counts** | **df** | **p-value** | **Estimate** | **95% CI** | **α** |
| --- | --- | --- | --- | --- | --- | --- |
| Cochran's Q | Q = 1.667 | 1 | 0.197 | — | — | 0.05 |
| Exact McNemar | 10 raw correct/normalized incorrect; 5 reverse; 15 discordant | — | 0.302 | — | — | 0.05 |
| Paired-bootstrap accuracy difference (raw − normalized) | — | — | — | 4.6 pp | −1.9 to 12.0 pp | 0.05 |

#### Appendix 10. Confidence across all out-of-fold test plants

Model confidence was assessed for all 108 out-of-fold predictions using the maximum predicted-class probability. Correct predictions had higher confidence than incorrect predictions (two-sided Mann–Whitney U = 995, p < 0.001; α = 0.05), indicating that confidence generally tracked prediction correctness. Errors occurred in all four experiments rather than being confined to one experiment. Maximum probabilities are interpreted as relative confidence scores rather than formally calibrated probabilities. Results are summarized in the tables below.

**Confidence according to prediction:**

| **Outcome or comparison** | **Plants** | **Percentage** | **Median maximum confidence** | **Statistic** | **Two-sided p** | **α** |
| --- | --- | --- | --- | --- | --- | --- |
| Correct | 96 | 88.9% | 0.953 | — | — | — |
| Incorrect | 12 | 11.1% | 0.677 | — | — | — |
| Correct versus incorrect | 108 | — | — | Mann–Whitney U = 995 | <0.001 | 0.05 |

**Incorrect predictions by experiment:**

| **Experiment** | **Test plants** | **Incorrect predictions** | **Error rate** |
| --- | --- | --- | --- |
| Experiment 1 | 18 | 1 | 5.6% |
| Experiment 2 | 30 | 5 | 16.7% |
| Experiment 3 | 30 | 4 | 13.3% |
| Experiment 4 | 30 | 2 | 6.7% |
| Total | 108 | 12 | 11.1% |

#### Appendix 11. Sensitivity to the two heat regimes

Experiment 1 (30/22°C) and Experiment 4 (30/27.5°C) heat regimes were compared after normalization to their respective within-experiment controls across 60 features (trait-by-time combinations). Their control-relative signatures were positively concordant, with 43/60 features (71.7%) changing in the same direction, although 22/60 features differed significantly between regimes in both Welch and Mann–Whitney tests after Benjamini–Hochberg correction (α = 0.05). Because both regimes imposed the same elevated daytime temperature and produced directionally consistent responses for most features, they were retained as a broad Heat class for distinguishing among major abiotic treatment categories and to maintain adequate class representation for model development. This grouping does not imply that the regimes were phenotypically equivalent, and the difference in night temperature remains a study limitation. Given the modest sample sizes, these results provide sensitivity evidence rather than a formal equivalence test. Results are summarized in the tables below.

**Sample sizes:**

| **Experiment and heat regime** | **Control plants** | **Heat plants** |
| --- | --- | --- |
| Experiment 1: 30/22°C | 9 | 9 |
| Experiment 4: 30/27.5°C | 4 | 6 |

**Control-relative signature comparison:**

| **Analysis** | **Estimate** | **Nominal p-value** | **Bootstrap 95% CI** | **Multiple-testing result** |
| --- | --- | --- | --- | --- |
| Pearson correlation | 0.626 | 8.98 × 10⁻⁸ | 0.332 to 0.696 | — |
| Spearman correlation | 0.596 | 4.95 × 10⁻⁷ | 0.374 to 0.737 | — |
| Cosine similarity | 0.710 | — | 0.449 to 0.766 | — |
| Effect-direction agreement | 71.7% | — | 58.3% to 83.3% | — |
| Welch feature tests | 22/60 significant | — | — | Benjamini–Hochberg q < 0.05 |
| Mann–Whitney feature tests | 22/60 significant | — | — | Benjamini–Hochberg q < 0.05 |
